# Dysregulation of a novel autophagosome-mitochondria contact contributes to autophagy dysfunction and neurodegeneration in tauopathy

**DOI:** 10.64898/2026.03.23.713823

**Authors:** Nuo Jia, Hongyuan Guan, Yantao Zuo, Yu Young Jeong, Niharika Amireddy, Gavesh Rajapaksha, Cuauhtemoc Ulises Gonzalez, Nora Jaber, Yun-Kyung Lee, Marialaina Nissenbaum, David J. Margolis, Wei Dai, Alexander W. Kusnecov, Qian Cai

## Abstract

Mitochondria engage in extensive communication with other organelles through membrane contacts. Perturbed mitochondria-organelle interactions are indicated in a variety of neurodegenerative diseases, but the underlying mechanisms remain poorly understood. Here, we report a new class of mitochondria-organelle communication: autophagosome/autophagic vacuole (AV)-mitochondria (Mito) contact, which exhibits hyper-tethering in tauopathy neurons, consequently hampering AV retrograde transport. Such defects are attributed to accelerated turnover of the contact release factor TBC1D15, triggered by mitochondrial bioenergetic deficit-induced hyperactivity of the AMP-activated protein kinase (AMPK). Increasing TBC1D15 levels or repressing AMPK activity normalizes AV-Mito contact release and restores retrograde transport of AVs, thereby increasing autophagic cargo clearance and reducing tau burden in tauopathy axons. Furthermore, overexpression of TBC1D15 enhances autophagic clearance and attenuates tau pathology, alleviating neurodegeneration and cognitive dysfunction in tauopathy mice. Taken together, our study provides new insights into AV-Mito contact dysregulation in tauopathy-related autophagy failure, laying the groundwork for the development of potential therapeutics to combat tauopathy diseases.

## Introduction

Mitochondria are cellular power plants that supply energy in the form of ATP to fuel various activities, essential for neuronal function and survival. Mitochondrial defects are a central concern in nervous system aging and are associated with major neurodegenerative disorders, including Alzheimer’s disease (AD), which is characterized by the presence of extracellular amyloid plaques consisting of amyloid β (Aβ) and intracellular neurofibrillary tangles (NFTs) composed of hyperphosphorylated tau (phospho-tau) in diseased brains (1, 2). Recent breakthroughs in the field make it increasingly evident that mitochondria interact with other organelles by forming membrane contacts. Such mitochondria-organelle communication is involved in the regulation of organelle dynamics and function (3–7). Disturbances in mitochondrial interactions with other organelles have been implicated in neurodegenerative diseases (5–7). For instance, abnormal mitochondria-lysosome contacts have been reported in Charcot-Marie-Tooth type 2 disease and Parkinson’s disease (8–10). In AD, advancing the understanding of how mitochondria communicate with other organelles remains a major goal. Some findings from early studies have revealed dysregulation of mitochondria-endoplasmic reticulum (ER) contacts that augments Aβ overproduction (11–13). However, whether and how dysregulation of mitochondria-organelle communication participates in tauopathy-related pathologies remains unanswered.

Autophagy constitutes a key mechanism of cellular quality control essential for neuronal health and maintenance. It has been established that pathological tau, a defining feature of AD and related tauopathies, is degraded within lysosomes after sequestration within autophagosomes/autophagic vacuoles (AVs) (14–17). Importantly, many studies have highlighted tau accumulation in tauopathy brains and pathological tau-induced neuronal toxicity (18–27). These findings imply that autophagy failure plays a pivotal role of in the buildup of pathological tau. Indeed, earlier work has uncovered aberrant accumulation of AVs in tauopathy patient brains (28), suggestive of autophagy defects. Studies in cultured neuronal cell lines have further demonstrated disturbance of autophagy function relevant to pathological tau expression (29). However, the mechanism underlying the link between autophagy failure and pathologic tau buildup remains elusive.

Mitochondrial defects are a fundamental early event in AD pathogenesis (30–33). In tauopathy, mitochondrial abnormalities have also been indicated (32, 34–37). Specifically, we and others have revealed that mitochondrial deficits and energy crisis occur early in tauopathy (38–41). Our recent work has further demonstrated that mitochondrial dysfunction precedes synaptic failure and tau pathology in tauopathy brains (41–43). However, whether and how such early mitochondrial deficiency contributes to the onset and progression of pathologies in tauopathy remains largely unknown. Notably, defects in complex I activity of the mitochondrial respiratory chain that led to metabolic disruption were found to correlate strongly with tau burden (44, 45), supporting a strong link between mitochondrial bioenergetic deficits and tau pathology in tauopathy. Given that mitochondrial dysfunction is an early feature of tauopathy, a fundamental question needs to be answered: Are such early mitochondrial deficits involved in the pathological accumulation of tau in tauopathy neurons?

The present study has unveiled a new class of inter-organelle interactions through AV-mitochondria (Mito) membrane contacts that display excessive contact tethering in tauopathy neurons. Such a defect results from a deficiency in AV-Mito contact release factor Rab7 GTPase activating protein (GAP) TBC1D15, attributable to hyperactivity of AMP-activated protein kinase (AMPK) triggered by mitochondrial energy metabolism dysfunction in tauopathy. As a result, AV-Mito contact release failure disrupts autophagy activity by halting retrograde axonal transport of AVs. Overexpression (OE) of TBC1D15 normalizes AV-Mito contact dynamics and reverses defective AV retrograde transport for pathologic tau clearance in axons, thereby ameliorating tau pathology and protecting against neurodegeneration and cognitive impairment in tauopathy mice. Thus, our findings provide new mechanistic insights into energy metabolism deficiency, and the resulting AV-Mito contact hyper-tethering in autophagy dysfunction in the context of tauopathy and have implications for understanding aging and other neurodegenerative diseases associated with mitochondrial and metabolic disruptions and autophagy dysfunction at large.

## Results

### Autophagosome/autophagic vacuole (AV)-mitochondria (Mito) contact hyper-tethering is coupled with AV retrograde transport defects in tauopathy axons

Given that autophagy abnormalities have been reported in the brains of tauopathy patients (28), we aimed to investigate their underlying mechanisms. We first performed Transmission Electron Microscopy (TEM) analysis of tauP301S Tg (PS19) mouse brains, focusing on presynaptic terminals, where AVs are primarily generated (46–51). Indeed, we found that AV-like structures were aberrantly accumulated at hippocampal presynaptic terminals in tauP301S Tg mice at 8 months of age, as evidenced by significant increases in the density of presynaptic AVs and the percentage of presynaptic terminals with AVs, suggestive of autophagic accumulation at tauopathy synapses (*Fig. 1A and B*). While this observation is in line with prior work (28), we also found that many of these AVs appeared to be in contact with mitochondria. Such AV-Mito contacts were not readily seen at the presynaptic terminals of non-Tg mouse brains. Moreover, AV-Mito contacts at tauP301S Tg synapses displayed reduced intermembrane distance and increased contact length between these two organelles, relative to those at non-Tg synapses (*Supplemental* Fig. S1 *B* and *C*). Of note, the average length and distance of these AV-Mito contact sites at tauopathy synapses are 135.32 nm ± 6.36 nm, and 16.49 nm ± 1.38 nm (mean ± SEM), respectively. To confirm this *in vivo* finding, we next characterized AV-Mito contact dynamics in the axons of live cortical neurons derived from tauP301S Tg mouse brains by carrying out high-resolution confocal time-lapse imaging. Prior to imaging, neurons were treated with the mTOR-independent autophagy inducer trehalose, which we and others previously showed to induce neuronal autophagy (15, 52, 53). Indeed, tauP301S Tg axons displayed marked increases in the frequency and duration of AV-Mito contact tethering, the number of AV-Mito contacts, and the percentage of AVs in AV-Mito contacts, compared to those in non-Tg axons (Fig. 1 *C-F*). It is worth noting that AVs were barely detectable in the dendrites of non-Tg and tauP301S Tg neurons, and thus, AV-Mito interactions were limited to the axons (*Supplemental* Fig. S1 *D* and *E*). In agreement with our TEM data, this observation indicates that AVs can form membrane contacts with mitochondria, whereas AV-Mito contact dynamics is likely impaired in tauopathy axons. To visualize these contact sites in three dimensions, we further conducted correlative light and electron microscopy (CLEM) and cryo-electron tomography (cryo-ET) analysis in tauP301S Tg cortical neuron culture. Using RFP-LC3 and MitoView Green to indicate AVs and mitochondria, respectively, CLEM enabled the identification of potential AV-Mito contact sites in the region of axonal processes for subsequent cryo-ET imaging. Tomograms collected at these sites revealed membrane contacts between the two organelles with intermembrane distances consistent with those observed in TEM analysis. Notably, densities bridging the AV and mitochondrial membranes were clearly observed, suggesting the presence of putative tethering complexes likely involved in AV-Mito contacts (Fig. 1 *G* and *Supplemental* Fig. S1 *F* and *G*). Combined, these results suggest that AV-Mito untethering is impaired in tauopathy axons, leading to autophagic accumulation by hampering AV retrograde transport.

**Figure 1.**
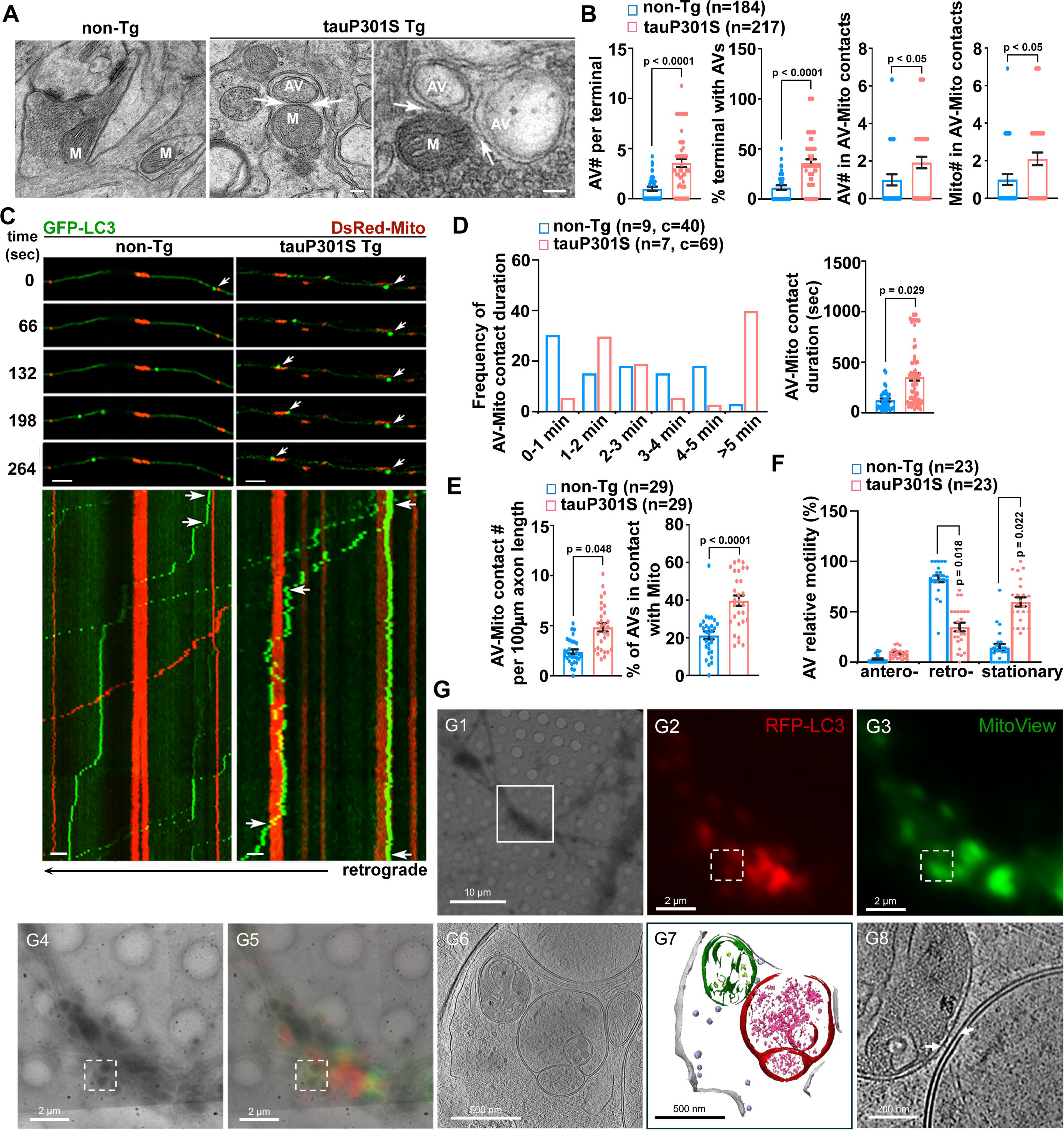
Autophagosome/autophagic vacuole (AV)-mitochondria (Mito) contact hyper-tethering and AV retrograde transport defects in tauopathy axons. (*A* and *B*) Representative transmission electron microscopy (TEM) images (*A*) and quantitative analysis (*B*) of 8-month-old tauP301S Tg (PS19) hippocampi. The number of presynaptic AVs, the percentage of terminals containing AVs, and the number of AVs or mitochondria in AV-Mito contacts were quantified and normalized to or compared to those in non-Tg littermate controls. Data were quantified from a total number of presynaptic terminals (*n*) as indicated in parentheses (*B*) from three mice per group. Arrows: AV-Mito contacts; AV: autophagic vacuole; M: mitochondrion. (*C*-*F*) Representative time-lapse images and kymographs (*C*) and quantitative analysis (*D*-*F*) of non-Tg and tauP301S Tg axons after 24-hour trehalose (100 mM) incubation. AV-Mito contact duration and its frequency, the contact number per 100 μm axonal length, AV percentage in contacts, and the relative motility of AVs were quantified and compared between the two groups, respectively. Data were collected from three dissections: the total numbers of neurons (*n*) and AV-Mito contacts (*c*) are indicated in parentheses (*D*-*F*). Arrows: AVs (time-lapse images in top *C*); AV-Mito contacts (kymographs in bottom *C*). (*G*) Correlative light and electron microscope (CLEM) and cryo-electron tomography (cryo-ET) analysis of AV-Mito contacts in tauopathy axons. Low-magnification cryo-EM image of a tauP301S Tg neuron showing a region of the axonal process (*G1*). CLEM of the region outlined by the white box in *G1*, showing fluorescence images of mRFP-LC3-labeled AVs (*G2*) and MitoView Green-marked mitochondria (*G3*). The cryo-EM image of the same region acquired at 3,600 × magnification (*G4*) is overlaid with the fluorescence image (*G5*). The white dashed box indicates the potential AV-Mito contact site selected for tilt-series acquisition. Slice view of the reconstructed tomogram (*G6*) with membrane annotations of a mitochondrion (green), an AV (red), synaptic vesicles (purple), and membranes (light grey) (*G7*). Zoomed-in view of the AV-Mito contact site showed densities bridging the two membranes (*G8*, white arrows). Data were expressed as the mean ± SEM and analyzed using linear mixed-effects models. Scale bars: 100 nm (*A*) and 10 μm (*C*).

### Rab7 GAP TBC1D15 deficiency causes AV-Mito contact hyper-tethering and AV retrograde transport defects in tauopathy axons

Next, we explored the mechanism underlying altered AV-Mito contact dynamics in tauopathy neurons. Our previous studies have established that newly generated AVs gain retrograde transport motility by being loaded with the dynein transport machinery through fusion with Rab7-positive late endosomes to form amphisomes (49). AVs in AV-Mito contacts might fail to fuse with late endosomes and then recruit the transport machinery needed for AV retrograde transport, thereby augmenting AV-Mito contact tethering. Given that amphisomes, but not nascent autophagosomes, are positive for Rab7, we thus determined whether Rab7 is associated with AVs in contact with mitochondria. We found that Rab7 was present on most AVs in tauopathy axons, including AVs in AV-Mito contacts (94.80% ± 1.3%) (Fig. 2 *A* and *B*). This result rules out the possibility that AV-Mito hyper-tethering and defective AV retrograde transport are the consequence of impaired AV-late endosome fusion and further indicates an alternative mechanism underlying AV-Mito contact release defects in tauopathy neurons. As a small GTPase that belongs to the Rab family, Rab7 regulates multiple intracellular trafficking pathways (54). We thus sought to address whether AV-Mito hyper-tethering could be attributed to the deregulation of Rab7 GTPase activity. We first treated tauP301S Tg neurons with CID 1067700, a competitive nucleotide binding inhibitor for Rab7 (55), which, indeed, corrected AV-Mito contact release defects, as evidenced by a decrease in the density of AV-Mito contacts in tauopathy axons (Fig. 2 *A* and *B*). Moreover, CID 1067700 treatment also led to the restoration of impaired AV retrograde transport in tauP301S Tg axons (*Supplemental* Fig. S2 *I* and *J*). As expected, we also observed similar rescue effects in tauopathy axons expressing Rab7A mutant T22N that harbors an inactive GTPase domain and acts as an inactive Rab7 (56), but not Rab7A or constitutively active Rab7A mutant Q67L (*Supplemental* Fig. S2 *A* and *B*). These results collectively suggest that excessive AV-Mito tethering and AV retrograde transport defects could be attributed to hyperactive Rab7 in tauopathy neurons.

**Figure 2.**
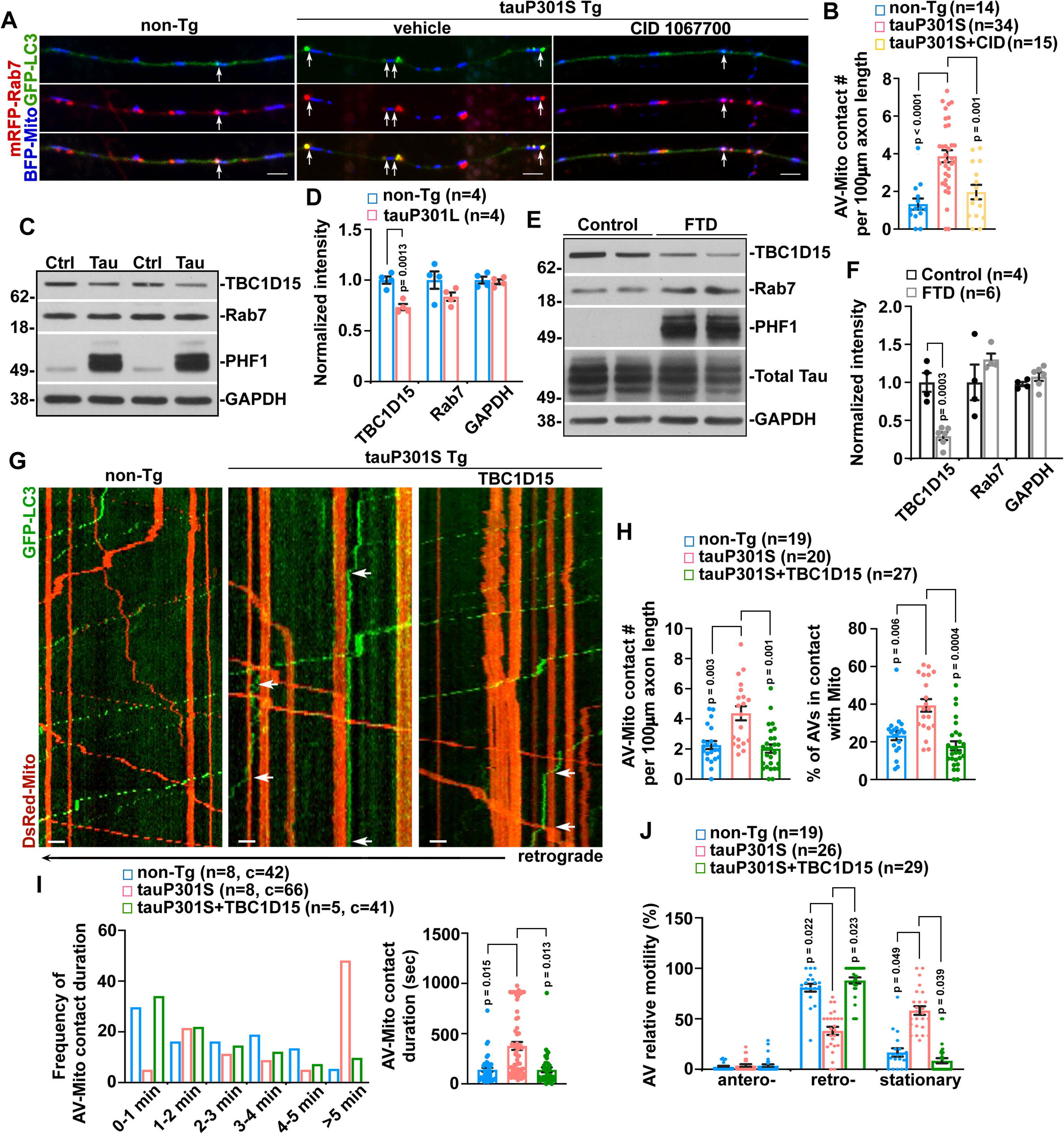
Rab7 GAP TBC1D15 deficiency causes excessive AV-Mito tethering and AV retrograde transport defects in tauopathy axons. (*A* and *B*) Representative images (*A*) and quantitative analysis (*B*) of AV-Mito contact tethering in non-Tg and tauP301S Tg axons after 24-hour trehalose treatment with and without CID 1067700 (80 nM). The data were expressed as AV-Mito contact number per 100 μm axonal length in non-Tg or tauP301S neurons. Data were collected from three dissections; the total numbers of neurons (*n*) are indicated in parentheses (*B*). Arrows: AV-Mito contacts. (*C*-*F*) Representative blots (*C* and *E*) and quantitative analysis (*D* and *F*) of protein levels in the brain of frontotemporal dementia (FTD) patients and 8-month-old tauP301L Tg (rTg4510) mice. Protein intensities were normalized to those in control human subjects or non-Tg mice. Data were quantified from four independent experiments. (*G*-*J*) Representative kymographs (*G*) and quantitative analysis (*H*-*J*) of AV-Mito contacts in tauP301S axons with TBC1D15 overexpression (OE). The data were quantified and expressed as AV-Mito contact duration and its frequency, the contact number per 100 μm axonal length, the percentage of AVs in contacts, and the relative motility of AVs in tauP301S axons with and without TBC1D15 OE, compared to those of non-Tg axons in the presence of trehalose (100 mM). Data were quantified from the total numbers of neurons (*n*) or AV-Mito contacts (*c*) as indicated in parentheses (*H*-*J*) from three dissections. Arrows: AV-Mito contacts. Data were expressed as the mean ± SEM and analyzed using linear mixed-effects models (*B* and *H*-*J*) or two-sided Student’s *t*-test (*D* and *F*). Scale bars: 10 μm.

TBC1D15 is known to function as a GAP for Rab7, facilitating Rab7 GTP hydrolysis and thereby inhibiting Rab7 activity (57, 58). Thus, we aimed to test whether hyperactive Rab7 might be due to TBC1D15 deficiency in tauopathy. We first examined TBC1D15 protein levels in the brains of tauP301L Tg (rTg4510) tauopathy mice and FTD patients. Indeed, despite no observed reductions in TBC1D15 mRNA levels in these tauopathy brains (*Supplemental* Fig. S2 *C* and *D*), TBC1D15 expression was significantly reduced in the brains of both tauopathy mice and FTD patients, while Rab7 showed no detectable changes (Fig. 2 *C*-*F*). These findings imply that TBC1D15 deficiency in Rab7 inhibition leads to Rab7 hyperactivity and consequently disrupts AV-Mito contact release in tauopathy neurons. We then assessed whether OE of TBC1D15 normalizes AV-Mito contact dynamics in tauP301S Tg axons. As expected, we found significant reductions in AV-Mito contact duration, AV-Mito contact density, and the percentage of AVs in AV-Mito contacts in tauP301S axons with TBC1D15 OE (Fig. 2 *G-I*). Moreover, defective AV retrograde movement was corrected in these tauopathy axons (Fig. 2 *J*). To further define the role of TBC1D15, we assessed AV-Mito contact dynamics in normal neurons with and without TBC1D15 RNAi. Neurons lacking TBC1D15 phenocopied defects in AV-Mito contact release in tauopathy neurons, as reflected by increases in AV-Mito contact duration and the percentage of AVs in contact with mitochondria and a decrease in AV retrograde transport (*Supplemental* Fig. S2 *E-H*), highlighting that TBC1D15 is required for maintaining AV-Mito contact dynamics and AV retrograde movement in axons. Combined, these findings support the notion that defects in AV-Mito contact release and the resulting AV retrograde transport impairment are attributed to TBC1D15 deficiency in tauopathy. Such defects could be a mechanism underlying autophagy dysfunction in tauopathy neurons.

### Mitochondrial bioenergetic stress-activated AMPK induces TBC1D15 deficiency by accelerating TBC1D15 turnover in tauopathy neurons

We next sought to determine whether mitochondrial deficits, an early feature of tauopathy (41, 42), contribute to TBC1D15 deficiency. In accord with the findings in tauopathy brains (Fig. 2 *C-F*), we found that the TBC1D15 level was lower in tauP301S Tg, relative to non-Tg neurons (Fig. 3 *A* and *Supplemental* Fig. S3 *A*). Furthermore, treatment with rotenone, a complex I inhibitor, resulted in additional TBC1D15 reduction in both non-Tg and tauP301S neurons. Similar effects were observed in neurons incubated with antimycin A, a complex III inhibitor (*Supplemental* Fig. S3 *B* and *C*). These observations imply that TBC1D15 decrease is likely a downstream effect of bioenergetic dysfunction. AMPK, a master energy sensor, is known to be activated upon metabolic stress within cells (59, 60). Indeed, previous work has reported AMPK overactivation in tauopathy brains (61). Consistently, we also observed increased AMPK activity in tauP301S Tg neurons, as reflected by the elevated ratio of pAMPKα/AMPKα (Fig. 3 *A* and *Supplemental* Fig. S3 *A*). As expected, rotenone treatment potentiated AMPK activity in both non-Tg and tauP301S neurons, suggesting that mitochondrial bioenergetic disruption effectively enhances AMPK activity. We then examined whether TBC1D15 deficiency is a consequence of AMPK activation. While increasing AMPK activity by 5-aminoimidazole-4-carboxamide riboside (AICAR), an analogue of AMP (62), exacerbated TBC1D15 deficiency in tauP301S neurons, such an effect was completely abrogated under co-treatment with Compound C (CC), an AMPK inhibitor (63), which led to even higher TBC1D15 levels than those under basal conditions (Fig. 3 *B* and *Supplemental* Fig. S3 *D*). This result indicates that the TBC1D15 reduction could be attributed to AMPK overactivation in tauopathy neurons.

**Figure 3.**
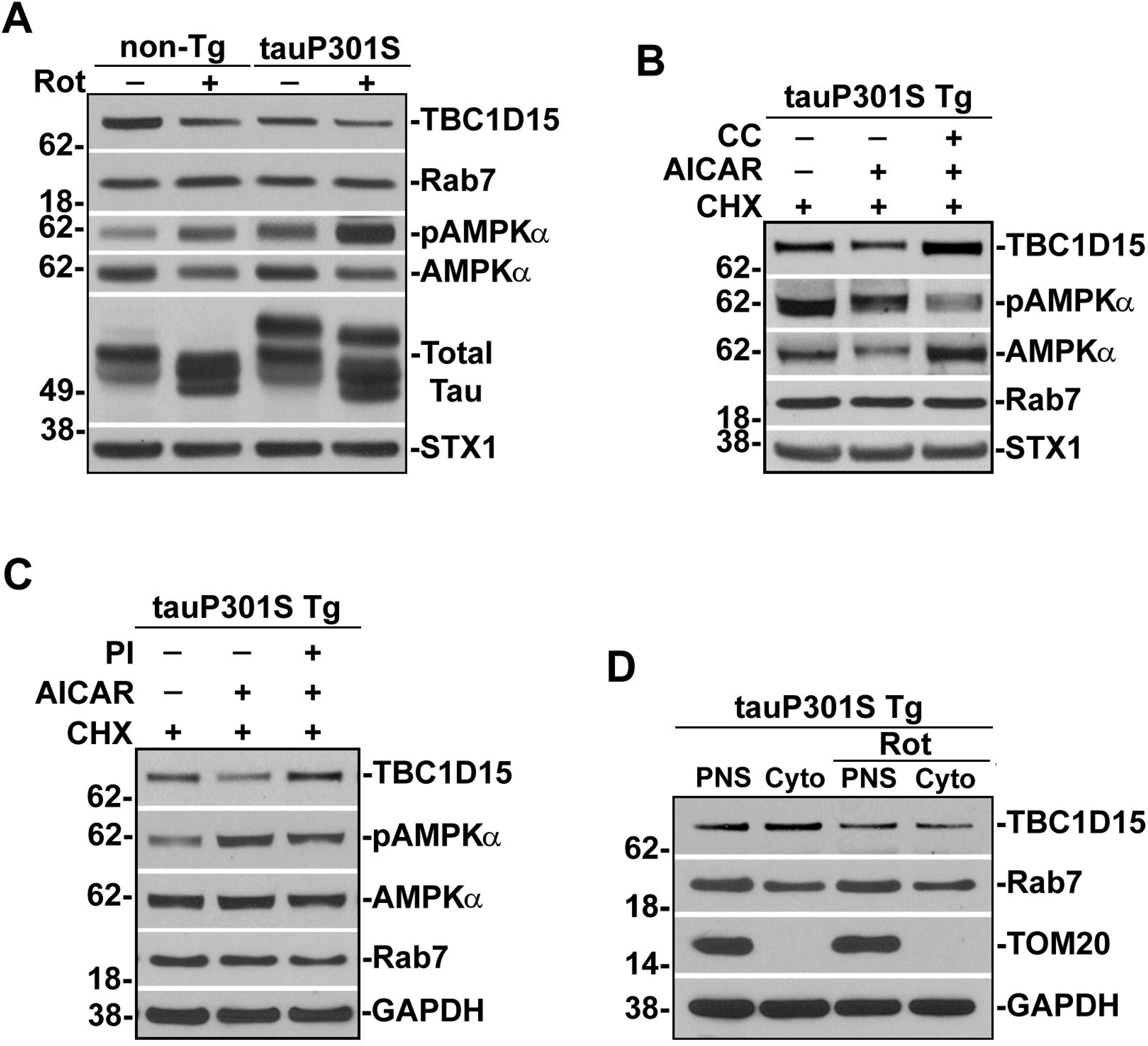
Bioenergetic stress-activated AMPK triggers TBC1D15 deficiency by accelerating TBC1D15 turnover in tauopathy neurons. (*A*) Representative blots of non-Tg and tauP301S Tg neurons with and without 24-hour rotenone (20 μM) incubation. Rot: rotenone. (*B*) Representative blots of tauP301S Tg neurons with 24-hour AICAR (0.5 mM) or/and CC (5 μM) incubation. CHX: cycloheximide (10 μg/ml); AICAR: 5-aminoimidazole-4-carboxamide riboside; CC: Compound C. (*C*) Representative blots of tauP301S neurons treated with and without AICAR and/or epoxomicin (100 nM). PI: proteasome inhibitor. (*D*) Representative blots of total and cytosolic TBC1D15 in tauP301S neurons with and without rotenone treatment. PNS: post-nuclear supernatant; Cyto: cytosolic fraction.

Tauopathy brains exhibited reduced levels of TBC1D15 protein but not mRNA (Fig. 2 *C-F* and *Supplemental* Fig. S2 *C* and *D*), suggestive of enhanced TBC1D15 turnover. In fact, AMPK signaling is known to promote the degradation activities of cellular components (59, 64–67). We found that inhibition of proteasome activity in tauP301S neurons effectively reversed AMPK-aggravated TBC1D15 reduction (Fig. 3 *C* and *Supplemental* Fig. S3 *E*). Similarly, in normal neurons, accelerated TBC1D15 turnover was detected upon AMPK activation, which was corrected in the presence of CC or epoxomicin, a proteasome inhibitor (*Supplemental* Fig. S3 *F* and *G*), suggesting a more general mechanism of AMPK-regulated TBC1D15 turnover. As the proteasome degrades cytosolic proteins (68, 69), rotenone treatment indeed led to a more pronounced reduction in the cytosolic pool of TBC1D15 (Fig. 3 *D* and *Supplemental* Fig. S3 *H*). These findings collectively indicate that AMPK hyperactivity augments TBC1D15 turnover through proteasome degradation in tau neurons. Additionally, autophagy induction showed no effects on TBC1D15 in tauP301S Tg neurons (*Supplemental* Fig. S3 *I* and *J*), excluding the possible involvement of autophagy in TBC1D15 deficiency. Together, these results support the view that AV-Mito contacts are controlled by AMPK, which senses cellular energy stress; bioenergetic dysfunction elevates AMPK activity, leading to enhanced TBC1D15 turnover and, consequently, defects in AV-Mito untethering in tauopathy neurons.

### AV-Mito contact release defects are attributed to hyperactivity of mitochondria-localized AMPK in tauopathy neurons

Mounting evidence has indicated compartmentalized regulation of AMPK signaling within cells (70, 71). In line with this pattern, mitochondria-localized AMPK activity has also been described (70, 72). Given that mitochondrial defects are an early feature in tauopathy (38–41), we next determined whether mitochondrial bioenergetic deficits have an impact on mitochondrial AMPK activity in neurons by applying Mito-ABKAR, a mitochondria-targeted Förster resonance energy transfer (FRET)-based AMPK biosensor (*Supplemental* Fig. S4 *A*) (70). We found that mitochondria in tauP301S axons exhibited an increase in FRET signal (YFP/CFP ratio) as compared to those in non-Tg neurons. Rotenone-induced bioenergetic disruption effectively augmented mitochondria-localized AMPK activity (*Supplemental* Fig. S4 *B* and *C*). To confirm our imaging data, we further purified mitochondria from tauP301S Tg neurons. Rotenone treatment led to a drastic increase in AMPK activity at these tauopathy mitochondria, as reflected by an elevation in the ratio of pAMPKα/AMPKα (*Supplemental* Fig. S4 *D* and *E*).

These findings prompted us to further determine whether the increase in mitochondria-localized AMPK activity results in AV-Mito contact hyper-tethering in tauopathy neurons. We utilized TOM20-mChF-AIP (Mito-AIP), a genetically encoded AMPK inhibitor peptide sequence subcloned into the mCherry-FKRP vector with the mitochondria-targeting sequence (MTS) of TOM20, which specifically inhibits mitochondrial localized AMPK activity (*Supplemental* Fig. S4 *F*) (70). Indeed, expressing Mito-AIP normalized AMPK activity at the mitochondria in tauP301S Tg axons, compared to that in non-Tg axons (*Supplemental* Fig. S4 *G* and *H*). Such a rescue effect was absent in control tauP301S Tg axons expressing Mito-AIP (TA), whose phosphorylation site was replaced with an Ala residue, leading to the abolition of its inhibitory effects on mitochondria-localized AMPK activity (*Supplemental* Fig. S4 *F*) (70). We then examined whether inhibition of mitochondrial AMPK activity is sufficient to correct AV-Mito contact release defects in tauopathy neurons. Strikingly, relative to control tauP301S axons expressing Mito-AIP (TA), Mito-AIP-expressing tauP301S axons displayed enhanced AV-Mito untethering, as reflected by drastic reductions in AV-Mito contact duration and numbers and the percentage of AVs in AV-Mito contacts. Such rescue effects were accompanied by a significant increase in AV retrograde transport (*Supplemental* Fig. S4 *I-L*). Besides, inhibition of AMPK activity at the lysosomes or Golgi failed to correct defects in AV-Mito untethering and AV retrograde transport in tauP301S Tg axons (*Supplemental* Fig. S5 *A-D*). Together, these findings suggest that mitochondrial bioenergetic deficits and the resulting hyperactivity of mitochondria-localized AMPK prevent AV-Mito untethering, thereby impeding AV retrograde transport in tauopathy axons.

### AV-bound TBC1D15 GAP activity mediates AV-Mito contact release and enables autophagic cargo removal from axons

We next performed multiple lines of imaging studies in normal neurons to elucidate the mechanism underlying TBC1D15-regulated AV-Mito contact dynamics. The primary localization of TBC1D15 at mitochondria was reported in non-neuronal cells (57, 58, 73). However, in neurons, the noncytosolic pool of TBC1D15 was found to be predominantly associated with AVs rather than mitochondria (Fig. 4 *A* and *B*). Importantly, these TBC1D15-targeted AVs moved in a predominantly retrograde direction along axons (Fig. 4 *C* and *D*), implying the likelihood that AV-bound TBC1D15 controls AV-Mito untethering. To exclude the possible involvement of mitochondria-localized TBC1D15, we assessed AV-Mito contact dynamics in neurons with loss of Fis1, the mitochondrial TBC1D15 anchor (57, 73). As expected, knocking down Fis1 led to no defects in AV-Mito contact release, as evidenced by no changes in AV-Mito contact duration relative to those in control neurons (*Supplemental* Fig. S6 *A-C*). Also, AV motility was unaltered in these axons (*Supplemental* Fig. S6 *D*). To further delineate the role of AV-bound TBC1D15, we conducted triple-channel time-lapse imaging in axons expressing TBC1D15 or TBC1D15 D397A mutant lacking TBC domain GAP activity (57, 74). While both TBC1D15 and TBC1D15 D397A were localized to AVs (Fig. 4 *E*), compared to control axons expressing TBC1D15, OE of TBC1D15 D397A mutant recapitulated tauopathy-linked AV-Mito contact release defects, resulting in marked increases in the duration and numbers of AV-Mito contacts and the percentage of AVs in AV-Mito contacts, accompanied by reduced AV retrograde transport (Fig. 4 *E-H*). Hence, these observations collectively indicate that AV-localized TBC1D15 untethers AV-Mito contacts through its GAP activity.

**Figure 4.**
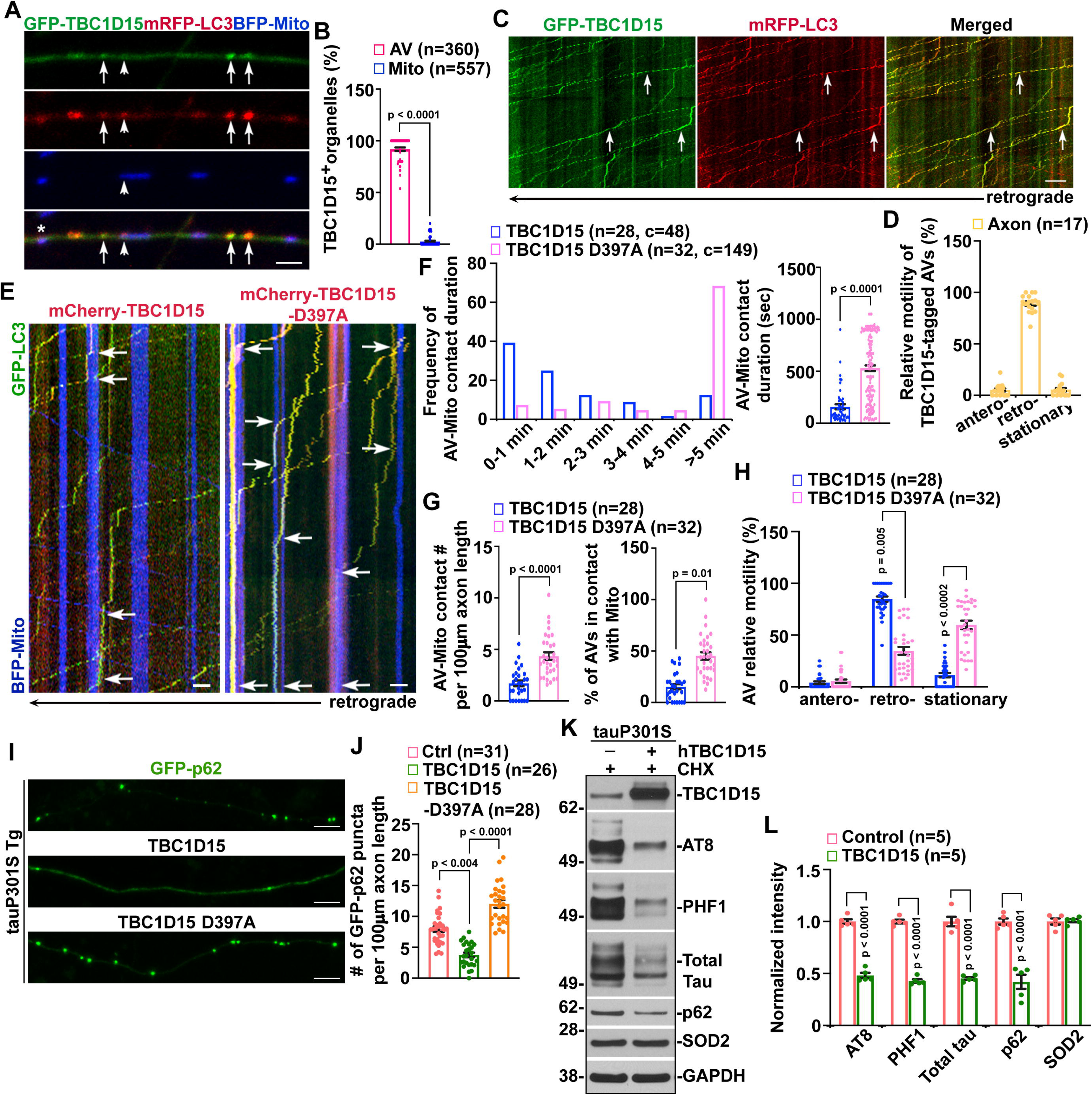
AV-Mito untethering relies on AV-bound TBC1D15 GAP activity and is required for autophagic cargo clearance in tauopathy axons. (*A* and *B*) Representative images (*A*) and quantitative analysis (*B*) of TBC1D15 localization in normal axons. Data were expressed as the percentage of AVs or mitochondria positive for TBC1D15 and quantified from the total numbers of AVs or mitochondria (*n*, indicated in parentheses) from the axons of 51 neurons (*B*). Arrows: TBC1D15-positive AVs; arrowheads: a TBC1D15-negative AV in an AV-Mito contact; asterisk: mitophagosome. (*C* and *D*) Representative kymographs (*C*) and quantitative analysis (*D*) of TBC1D15-tagged AVs in axons with 24-hour trehalose incubation. The relative motility of AVs was quantified from the axons of 17 neurons as indicated in parentheses (*D*). Arrows: TBC1D15-tagged motile AVs. (*E*-*H*) Representative kymographs (*E*) and quantitative analysis (*F*-*H*) of AV-Mito contacts in axons expressing TBC1D15 or TBC1D15 D397A mutant. The data were quantified and expressed as AV-Mito contact duration and its frequency, the contact number per 100 μm axonal length, the percentage of AVs in contacts, and the relative motility of AVs. Data were quantified from the total numbers of neurons (*n*) and AV-Mito contacts (c) as indicated in parentheses (*F*-*H*). Arrows: AV-Mito contacts. (*I* and *J*) Representative images (*I*) and quantitative analysis (*J*) of p62 in tauP301S axons with and without OE of TBC1D15 or TBC1D15 D397A mutant. The data were quantified and expressed as GFP-p62 puncta number per 100 μm axonal length. Data were quantified from the total numbers of neurons (*n*) as indicated in parentheses (*J*). (*K* and *L*) Representative blots (*K*) and quantitative analysis (*L*) of tauP301S Tg neurons transduced with and without AAV-hTBC1D15. Protein intensities were normalized to those in control tauP301S neurons without TBC1D15 OE. Data were collected from two (*B* and *D*), three (*F-H* and *J*), or five (*L*) dissections. Data were expressed as the mean ± SEM and analyzed using linear mixed-effects models (*B*, *F*-*H*, and *J*) or by two-sided Student’s *t*-test (*L*). Scale bars: 10 μm.

Given that AV retrograde transport returns autophagic cargos to the soma for lysosomal clearance (46–51, 75), we thus determined the effects of TBC1D15-mediated AV-Mito untethering on autophagic cargo clearance in axons. We first assessed p62/SQSTM1, an autophagy substrate, in tauopathy axons. While p62 punctate-like structures were accumulated in distal tauP301S Tg axons, OE of TBC1D15 led to a significant decrease in p62 accumulation (Fig. 4 *I* and *J*). Such a rescue effect was absent with OE of TBC1D15 D397A. Consistently, in normal neurons, autophagy induction reduced p62 puncta, but such an effect was reversed in axons with OE of TBC1D15 D397A (*Supplemental* Fig. S6 *E* and *F*). These findings suggest that disruption of TBC1D15-mediated AV-Mito untethering can lead to axonal accumulation of autophagic cargos. Phospho-tau is known as an autophagy substrate targeted for lysosomal degradation (14–17). Thus, we examined phospho-tau accumulation in tauP301S Tg axons and found that TBC1D15 OE reduced phospho-tau levels, as revealed by AT8 (S202/T205) and PHF1 (S396/S404) antibodies (*Supplemental* Fig. S6 *G*-*J*). To corroborate our imaging data, we further biochemically assessed tau levels in tauP301S Tg neurons after AAV-hTBC1D15 transduction. Consistent with our imaging data, TBC1D15 OE in tauP301S neurons decreased phospho-tau, as revealed by AT8 and PHF1 antibodies, and total tau, as visualized by Tau5 antibody, accompanied by p62 reduction (Fig. 4 *K* and *L*). Combined with the rescue effects of CID 1067700 treatment or TBC1D15 OE on AV-Mito contact release defects and AV retrograde transport impairment (Fig. 2 *A* and *B*, *G-J* and *Supplemental* Fig. S2 *I* and *J*), these observations suggest that rectifying AV-Mito hyper-tethering restores autophagy activity for pathological tau clearance in tau neurons. Taken together, these results highlight the critical role of GAP TBC1D15 in governing AV-Mito untethering and neuronal autophagy, further supporting the notion that AV-Mito contact defects contribute to tauopathy-associated autophagy dysfunction.

### Increasing TBC1D15 levels rectifies AV-Mito untethering defects and reduces p62 accumulation and tau pathology in tauopathy mouse brains

The *in vitro* rescue effects of TBC1D15 OE in cultured tauopathy neurons prompted us to determine its *in vivo* effects in tauopathy mouse brains by bilateral injection of AAV-mCherry-hTBC1D15 into the hippocampus and the cortex, an *in vivo* delivery procedure established by us and others (75–81). The injected mouse brains exhibited widespread mCherry fluorescence signals (*Supplemental* Fig. S7 *A* and *B*). The signals were also observable in the soma and processes of hippocampal neurons and cortical neurons of tauP301S Tg mouse brains. We first performed TEM analysis of AV-Mito contacts at presynaptic terminals in tauP301S Tg hippocampi injected with AAV-mCherry or AAV-mCherry-hTBC1D15. Compared to AAV-mCherry-injected control tauP301S mice, OE of TBC1D15 via AAV-mCherry-hTBC1D15 injection led to significant decreases in AV accumulation and the number of AVs or mitochondria in AV-Mito contacts at tauP301S Tg synapses (Fig. 5 *A* and *B*). Moreover, the AV-Mito contact intermembrane distance was increased, coupled with reduced contact lengths in these tau brains (*Supplemental* Fig. S7 *C* and *D*). These TEM data suggest that TBC1D15 OE-induced AV-Mito untethering decreases synaptic autophagic accumulation through enhancing AV retrograde transport.

**Figure 5.**
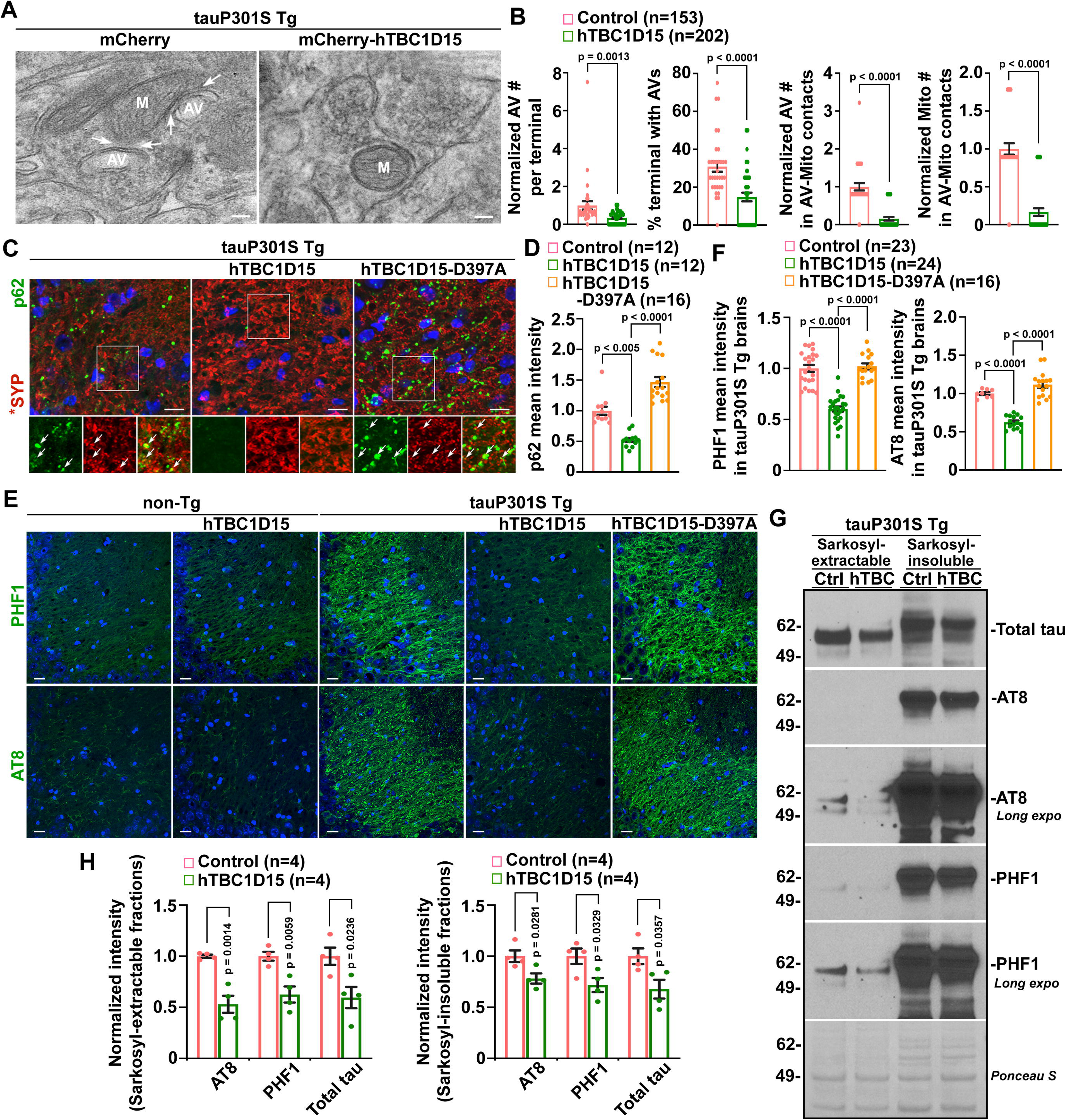
TBC1D15 OE reduces AV-Mito hyper-tethering and alleviates p62 accumulation and tau pathology in tauopathy mouse brains. (*A* and *B*) Representative TEM images (*A*) and quantitative analysis (*B*) of the hippocampi in 8-month-old tauP301S Tg mouse brains infected with AAV-mCherry or AAV-mCherry-hTBC1D15. The number of presynaptic AVs, the percentage of presynaptic terminals containing AVs, and the number of AVs or mitochondria in AV-Mito contacts were quantified and normalized to or compared to those in control tauP301S Tg mouse brains. Data were collected from two mice per group; the total numbers of presynaptic terminals (*n*) are indicated in parentheses (*B*). (*C* and *D*) Representative images (*C*) and quantitative analysis (*D*) of p62 accumulation in 10-month-old tauP301S Tg mouse hippocampi with and without OE of TBC1D15 or TBC1D15 D397A. p62 fluorescence mean intensity in the hippocampal mossy fibers was quantified and normalized to that of AAV-mCherry-injected tauP301S Tg controls. Data were collected from three mice per group; the total numbers of imaging slice sections (*n*) are indicated in parentheses (*D*). SYP: synaptophysin. Arrows: p62 clusters at SYP-indicated presynaptic terminals. (*E* and *F*) Representative images (*E*) and quantitative analysis (*F*) of tau accumulation in 10-month-old non-Tg and tauP301S Tg hippocampi with and without OE of TBC1D15 or TBC1D15 D397A. AT8 and PHF1 antibody-marked phospho-tau mean intensities in the hippocampal mossy fibers were quantified and normalized to those of AAV-mCherry-injected tauP301S Tg control mice. Data were collected from three mice per group; the total numbers of imaging slice sections (*n*) are indicated in parentheses (*F*). (*G* and *H*) Representative blots (*G*) and quantitative analysis (*H*) of AT8 and PHF1 antibody-marked phospho-tau and Tau5 antibody-indicated total tau in sarkosyl-extractable and sarkosyl-insoluble fractions from 10-month-old tauP301S Tg mouse brains with and without TBC1D15 OE. Protein intensities were normalized to those in AAV-mCherry-injected tauP301S Tg controls. Data were quantified from four mice per group. Data were expressed as the mean ± SEM and analyzed using linear mixed-effects models (*B, D* and *F*) or by two-sided Student’s *t*-test (*H*). Scale bars: 100 nm (*A*), 10 μm (*C*), and 20 μm (*E*).

We next performed immunohistochemistry and found that p62 was clustered in the hippocampal mossy fibers of tauP301S Tg mice, with a marked localization at presynaptic terminals, as indicated by synaptophysin (SYP). TBC1D15 OE in tauP301S Tg mouse brains induced a robust reduction in p62, and such rescue effects were absent in AAV-TBC1D15 D397A-injected tauP301S Tg mouse brains (Fig. 5 *C* and *D*). In non-Tg mouse brains, p62 clusters were undetectable regardless of TBC1D15 OE (*Supplemental* Fig. S7 *E*). This finding is consistent with our imaging data from cultured tau neurons (Fig. 4 *I* and *J*), suggesting that TBC1D15-mediated rescue effects are attributed to its GAP activity. We then performed additional biochemical analyses in these AAV-injected mouse brains. Consistent with results from our *in vitro* studies and *in vivo* imaging data (Fig. 4 *I*-*L* and Fig. 5 *C* and *D*), p62 levels were decreased in tauP301STg mouse brains with TBC1D15 OE (*Supplemental* Fig. S7 *F* and *G*). It is worth noting that these changes were accompanied by increases in synaptic proteins, including SYP and SNAP25, suggesting TBC1D15 OE-induced neuroprotective effects. As expected, we did not detect significant TBC1D15 OE-associated changes in the levels of p62 along with SYP and SNAP25 in non-Tg mouse brains.

Given the robust phospho-tau reduction in cultured tau neurons with TBC1D15 OE (Fig. 4 *K* and *L* and *Supplemental* Fig. S6 *G-J*), we next determined the impact of TBC1D15-enhanced AV-Mito untethering on tau pathology in tauP301S Tg mouse brains. As expected, TBC1D15 OE led to drastic reductions in tau pathology as revealed by decreased phospho-tau levels in the hippocampus of tauP301S Tg mice, relative to those in control tauP301S mice injected with AAV-mCherry or AAV-mCherry-TBC1D15 D397A (Fig. 5 *E* and *F*). Of note, immunostaining with AT8 and PHF1 antibodies did not reveal detectable TBC1D15 OE-associated changes in the hippocampal phospho-tau of non-Tg mice. We further examined TBC1D15 OE-induced tau changes by employing a standard sarkosyl extraction protocol for investigating insoluble tau aggregates in brains (82–85). Previous studies have shown that sarkosyl-insoluble tau displays the same structural and antigenic properties as paired helical filaments (PHFs) isolated from NFTs and is distinguishable from normal, soluble tau proteins (86). Consistent with prior work, we found that sarkosyl-insoluble fractions were enriched with tau with higher molecular weight (above 62 KDa), in contrast to the presence of normally sized 50-60 KDa tau in sarkosyl-extractable fractions (83–85). More importantly, relative to control tauP301S Tg mice, TBC1D15 OE led to decreases in phospho-tau levels in both sarkosyl-extractable and sarkosyl-insoluble fractions (Fig. 5 *G* and *H*), which is consistent with our *in vitro* imaging and biochemical observations along with *in vivo* imaging data (*Supplemental* Fig. S6 *G-J*, Fig. 4 *K* and *L,* and Fig. 5 *E* and *F*). Therefore, our results support the view that TBC1D15 OE and the resulting AV-Mito untethering promote autophagy for tau clearance, thereby preventing the buildup of pathological tau in tauopathy brains.

### TBC1D15-enhanced AV-Mito contact release attenuates synapse loss and mitigates neuronal death in tauopathy mouse brains

Given evidence of tau pathology attenuations after TBC1D15 OE (Fig. 5 *E*-*H*), we further determined whether its enhancement of AV-Mito contact release and the resulting autophagic clearance could produce neuroprotective effects in tauopathy mouse brains. Compared to non-Tg littermate mice, we found that the mean intensity of SYP-indicated presynaptic terminals was significantly reduced in the hippocampal mossy fiber regions of tauP301S Tg mice (Fig. 6 *A* and *B*), suggesting synapse loss in these tauopathy mouse brains. Strikingly, such a defect is attenuated in tauP301S Tg mouse brains injected with AAV-mCherry-TBC1D15, but not AAV-mCherry-TBC1D15 D397A. This result indicates that the rescue effects of TBC1D15 GAP activity on tauopathy-linked synaptic degeneration are very likely attributed to enhanced autophagic clearance of phospho-tau. We further examined whether increasing TBC1D15 expression has a beneficial effect against neuronal death in tauopathy mouse brains. In accord with previous studies from other groups (18, 77), our data showed that tauP301S Tg mice exhibited decreased neuron density in the hippocampal CA3 regions relative to that in non-Tg littermate controls (Fig. 6 *A* and *B*). Importantly, TBC1D15 OE prevented neuronal loss, as evidenced by an increased number of NeuN-positive neurons in this region of tauP301S mouse brains. We did not observe such rescue effects in tauP301S Tg mouse brains injected with AAV-mCherry-TBC1D15 D397A, nor did we detect a significant change in the densities of presynaptic terminals and NeuN-positive neurons in the hippocampus of non-Tg mouse brains with TBC1D15 OE (Fig. 6 *A* and *B*). Together, our *in vivo* evidence from tauopathy mouse brains suggests that TBC1D15-enhanced AV-Mito contact release prevents tauopathy-associated synaptic defects and neurodegeneration.

**Figure 6.**
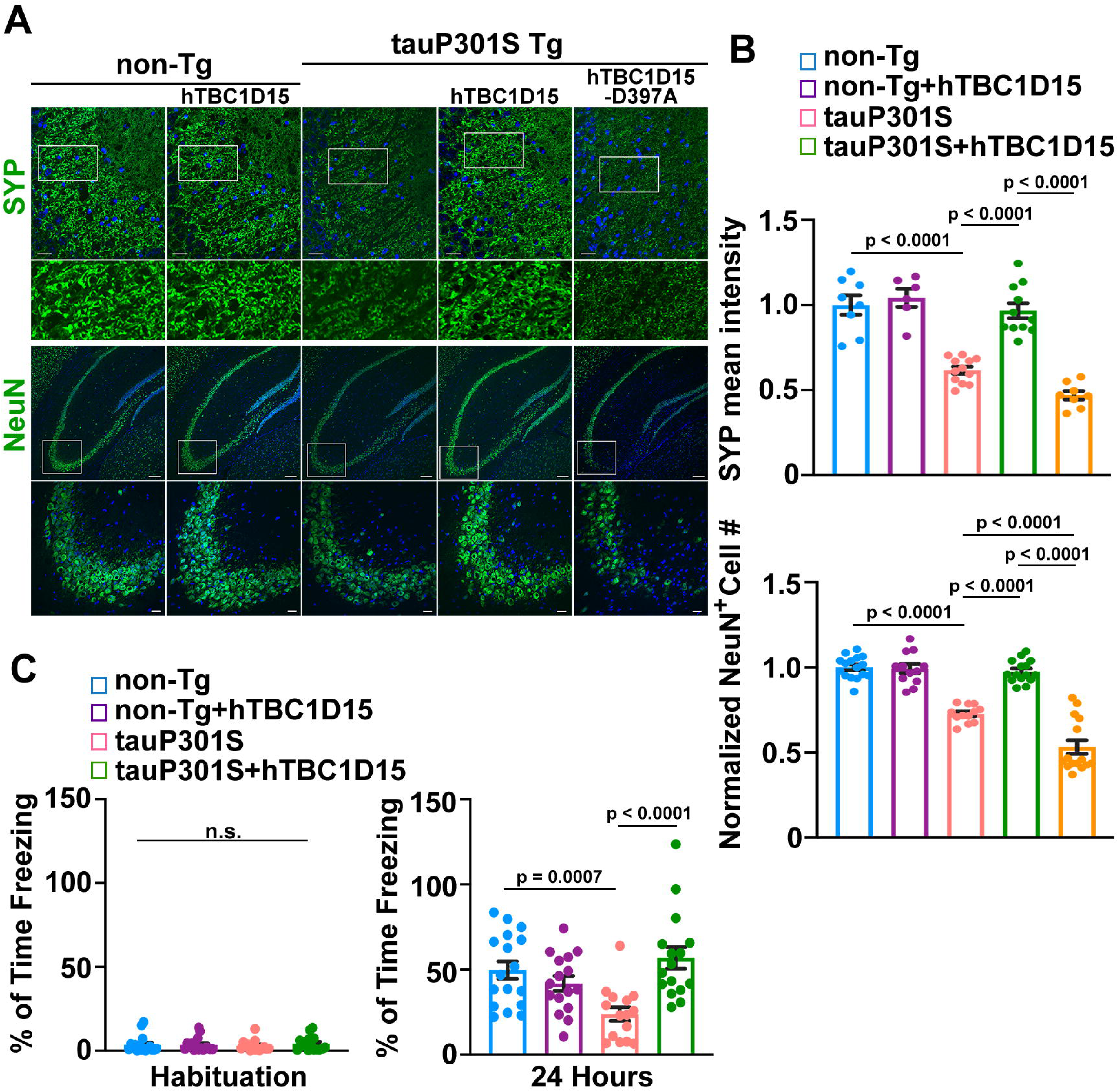
Attenuation of neurodegeneration and memory deficits in tauopathy mice with TBC1D15 OE. (A and *B*) Representative images (*A*) and quantitative analysis (*B*) of presynaptic terminal and neuron densities in 10-month-old non-Tg and tauP301S Tg hippocampi with and without OE of TBC1D15 or TBC1D15 D397A. The mean intensity of SYP-indicated presynaptic terminals in the hippocampal mossy fibers and the number of NeuN-labeled neurons in hippocampal CA3 areas marked by rectangles (Zoomed-in images shown in the lower row of each panel in *A*) were quantified and normalized to those of AAV-mCherry-injected non-Tg control mice. Data were collected from three mice per group. (*C*) Contextual fear conditioning task performed in 7-month-old non-Tg and tauP301S Tg male mice infected with AAV-mCherry or AAV-mCherry-hTBC1D15 (*n* = 15-17 male mice per group). Data were expressed as the mean ± SEM and analyzed using linear mixed-effects models (*B)* or by one-way ANOVA, followed by Sidak’s post-hoc test (*C*). Scale bars (*A)*: 20 μm (upper row, SYP), 250 μm (upper row, NeuN), and 25 μm (lower row, NeuN).

### Elevated TBC1D15 expression improves behavioral performance in tauopathy mice

We further determined whether increasing TBC1D15 expression ameliorates tauopathy-related learning and memory deficits, which are readily detectable in tauP301S Tg mice. We first examined sociability and social recognition memory in non-Tg and tauP301S Tg mice by conducting the three-chamber social investigation test (87–90). This test was conducted in two phases. Phase 1 assessed sociability to a stranger mouse (S1), while in the second phase, test subjects were allowed to explore the same mouse (S1) and a new stranger mouse (S2). For phase 1, animals with normal sociability display a preference for the S1-containing chamber over an empty chamber and are shown as having a longer time spent in the S1 chamber. For this phase, tauP301S Tg mice exhibited social interaction comparable to non-Tg littermate controls (*Supplemental* Fig. S8 *A*). However, during Phase 2, the social novelty preference/recognition test, which requires hippocampal function, non-Tg mice spent significantly more time exploring the chamber with a novel, stranger mouse (S2), while tauP301S Tg mice failed to distinguish the S2 stranger mouse from the familiar mouse (S1), spending similar amounts of time with both social targets (p = 0.8933). This result is in agreement with prior work by us and others (41, 42, 90), suggesting a social memory impairment in these tauP301S Tg mice. Interestingly, elevating TBC1D15 levels in tauP301S Tg mice rescued this phenotype, as evidenced by more time spent with the S2 stranger mouse (p = 0.0362). These observations suggest that TBC1D15 OE-enhanced AV-Mito contact release and AV clearance alleviate tauopathy-linked social recognition memory defects.

We next performed the Morris water maze (MWM) test to investigate spatial learning and memory in 7-month-old non-Tg and tauP301S Tg mice injected with AAV-mCherry-TBC1D15 or AAV-mCherry control. Mice were trained to find a hidden platform across four training blocks (Blocks 1-4) on a single day, with each block consisting of 3 to 4 acquisition trials. At one hour and 24 hours after the last hidden-platform training, memory testing for the location of the hidden platform was conducted in a 60-second probe trial (91, 92). Non-Tg mice with or without TBC1D15 OE showed similar latencies to find the platform during the hidden-platform training phase and favored the target quadrant in the probe trial (*Supplemental* Fig. S8 *B-D*), indicating that TBC1D15 OE had no beneficial or adverse effects on learning and memory in non-Tg mice. In contrast, tauP301S Tg mice without TBC1D15 OE performed poorly in both the acquisition phase and the memory probe tests, as reflected by longer latencies on training blocks 2-4 and less path length and less time in the target quadrant during probe trials that were conducted at one hour or 24 hours after training trials (p < 0.01) (non-Tg vs. tauP301S Tg). Importantly, AAV-mediated TBC1D15 OE in tauP301S Tg mice led to significantly improved learning and memory retention, to the extent that these animals displayed no detectable difference from non-Tg mice in latency and time in the target quadrant during the acquisition and probe trials, respectively (*Supplemental* Fig. S8 *B-D*). Combined, our results indicate that TBC1D15 elevation attenuates deficits in spatial learning and memory in tauP301S Tg mice.

We further examined the effects of TBC1D15 OE at 7 months of age on hippocampal-dependent memory in a contextual-based fear conditioning task, using freezing behavior as an index of fear-associated contextual memory (93). Freezing in tauP301S Tg mice injected with AAV-mCherry or AAV-mCherry-TBC1D15 did not differ from non-Tg littermates during the habituation phase that preceded footshock (Fig. 6 *C*). However, subsequent to the footshock exposure, non-Tg mice with AAV-mCherry or AAV-mCherry-TBC1D15 injection showed pronounced contextual fear memory, with 49.68% and 41.86% of the time spent “freezing” in the operant chamber. In contrast, tauP301S Tg mice spent only 23.84% of the time freezing (p = 0.0007) (non-Tg vs. tauP301S Tg). This observation is consistent with previous studies (41, 42, 90, 94), indicating a hippocampus-dependent memory deficit in this tauopathy model. Moreover, relative to control tauP301S Tg mice, TBC1D15 OE in this strain led to protective effects against the learning deficit, to the extent that the percent freezing time was similar to non-Tg mice. These findings support the view that TBC1D15-enhanced AV-Mito contact release at tauopathy synapses protects against tauopathy-related hippocampal-dependent memory impairment.

## Discussion

An enormous body of work has shown that membrane-enclosed organelles serve as interconnected hubs and interact with each other extensively through membrane contacts. Moreover, disturbances of inter-organelle interactions drive multi-organelle dysfunction and have thus been proposed as an emerging mechanism underlying aging and neurodegenerative diseases. Mitochondria dynamically form contacts with other organelles, and mitochondria-organelle communication has been shown to regulate organelle dynamics and functions (3–6). Some prior work has demonstrated aberrant ER-mitochondrial contacts and their participation in AD pathogenesis (11–13). Given that mitochondrial abnormalities are a characteristic feature of early tauopathy (38–41), two fundamental questions exist as to whether inter-organelle communication between mitochondria and other organelles is dysregulated in tauopathy and whether such a mechanism plays a role in the pathogenesis of tauopathy.

The current study has revealed, for the first time, a new class of inter-organelle interactions, AV-Mito contacts, which show untethering defects in tauopathy neurons, which is caused by reduced expression of TBC1D15, the contact release factor. We provided multiple lines of *in vitro* and *in vivo* evidence showing that, in cultured tauopathy neurons, defects in AV-Mito untethering hinder AV retrograde transport, thus disrupting autophagy activity for phospho-tau clearance. Conversely, increasing TBC1D15 levels rescues retrograde movement of AVs by rectifying AV-Mito contact dynamics and leads to drastic reductions in autophagic accumulation and phospho-tau levels in tau axons (Fig. 2 *J* and Fig. 5 *I-L*). TBC1D15 OE in tauopathy mouse brains corrects AV-Mito contact hyper-tethering and enhances autophagic clearance, which attenuates tau pathology and alleviates memory deficits in these mice. Collectively, these findings highlight a pivotal role of AV-Mito contact dysregulation in autophagy dysfunction, which leads to the buildup of pathological tau and tau-mediated neurotoxicity in tauopathy (*Supplemental* Fig. S9).

Excessive AV-Mito contact tethering in tauopathy neurons is the consequence of deficiency of the contact release factor TBC1D15, which can be rescued by inhibition of Rab7 activity or TBC1D15 OE. Notably, several recent studies have shown that Rab7 GTP hydrolysis via TBC1D15 also untethers mitochondria-lysosome contacts (8, 58, 95). Together with the current findings, these data suggest that a common molecular machinery is shared between mitochondrial interactions with AVs and with lysosomes. Nevertheless, it is worth noting that AV-Mito contacts exhibit several unique features. First, unlike mitochondria-lysosome contacts that are present in all distinct compartments of neurons (8), AV-Mito contacts are mainly observed in axons but are scarcely seen in dendrites (Fig. 1 and *Supplemental* Fig. S1 *D* and *E*), which is consistent with vigorous AV biogenesis occurring in distal axons, but not in dendrites (96–99). Second, AV-Mito contacts in normal axons occur more frequently with a relatively longer duration (Density (# per 100 μm axon length): 2.63 ± 0.42; Duration: 133.2 ± 17.10 s), compared to previously described mitochondria-lysosome contacts (Density: 0.93 ± 0.04; Duration: 112.5 ± 13.2 s) (8). Moreover, relative to normal axons, AV-Mito contacts occur more frequently with a markedly prolonged tethering duration in tauopathy axons (Density: 5.27 ± 0.96; Duration: 389.21 ± 36.00 s) (Fig. 1 *D* and *E*). Third, most Rab7-associated organelles are AVs in tauopathy axons (94.80% ± 1.3%) (Fig. 2 *A* and *B*), indicating that AV-Mito contact defects are a key phenotype attributed to Rab7 hyperactivity. Indeed, AV-Mito hyper-tethering and AV retrograde transport defects can be reversed by Rab7 inhibition or TBC1D15 OE (Fig. 2 *A* and *B*, *G-J* and *Supplemental* Fig. S2 *A* and *B*). Fourth, while mitochondria-lysosome contact release is shown to be mediated by mitochondria-bound TBC1D15 (8, 58), AV-Mito untethering is controlled by non-mitochondrial TBC1D15 that is most likely recruited to AVs from the cytosol (Fig. 4 *A-D* and *Supplemental* Fig. S6 *A-D*). GAP activity of AV-bound TBC1D15 is required for the untethering (Fig. 4 *E-H*). Therefore, these results define a new class of inter-organelle contacts in neurons and suggest that AV-Mito contact defects are a unique characteristic phenotype of tauopathy neurons.

In line with previous work in non-neuronal cell lines showing that TBC1D15 turnover is mediated by the proteasome system (8), we have further shown that, in neurons, such a mechanism is controlled by AMPK activity in response to the changes in mitochondrial bioenergetic status (Fig. 3 and *Supplemental* Fig. S3). Mitochondrial bioenergetic disruption in tauopathy neurons triggers the depletion of TBC1D15 in the cytosol where the proteasome is abundant (Fig. 3 *D* and *Supplemental* Fig. S3 *H*). AMPK activity localized at the mitochondria has been indicated previously (70, 72). Moreover, an early study reported that exercise-induced mitochondrial energy metabolic stress stimulates AMPK activity located at the outer mitochondrial membrane (OMM) (72). In accord with this study, our findings further demonstrate that mitochondria-localized AMPK responds effectively to mitochondrial bioenergetic deficits and is hyperactive in tauopathy neurons (*Supplemental* Fig. S4 *B-E*). Furthermore, such mitochondria-localized AMPK disrupts AV-Mito contact untethering, thus impeding AV retrograde transport in tauopathy axons (*Supplemental* Fig. S4 *I-L*). These findings raise a question about how mitochondria-localized AMPK activity regulates TBC1D15 turnover in the cytosol. Besides compartmentalized regulation of AMPK, a recent study proposed that spatial regulation of the AMPK signaling mechanism enables compartmentalized AMPK to be coordinated within cells (71). Thus, in tauopathy neurons, bioenergetic deficits sensed by mitochondrial AMPK likely trigger a mechanism that enables mitochondria-to-cytosol AMPK signaling, thereby stimulating proteasomal activity to degrade TBC1D15. Additionally, mitochondria-localized AMPK activity might directly enhance the proteasome-mediated turnover of TBC1D15 in the cytosol (*Supplemental* Fig. S9). Also, previous work reported increased proteasome recruitment to stressed mitochondria, which led to the degradation of multiple OMM proteins (100, 101). Excessive AV-Mito tethering might, in turn, facilitate the degradation of AV-bound TBC1D15 by the proteasome associated with mitochondria.

In non-neuronal cells, TBC1D15 was previously shown to localize primarily to mitochondrial membranes, where it participates in mitophagy and ER-curated mitochondrial fission (57, 58, 73). However, we did not detect a significant association of TBC1D15 with mitochondria in neurons with induced autophagy (Fig. 4 *A* and *B*). Knocking down Fis1, the mitochondrial TBC1D15 anchor (57, 73), showed no effects on AV-Mito contact dynamics and AV retrograde movement in axons (*Supplemental* Fig. S6 *A-D*). These results collectively suggest that mitochondria-bound TBC1D15 is not involved in regulating AV-Mito contacts in neurons, suggesting that alternative mechanisms mediate TBC1D15 delivery to AVs. Indeed, studies have shown that, owing to its multiple LC3-interaction region motifs, TBC1D15 interacts with LC3 and other ATG8 proteins (102, 103). Given that TBC1D15 is abundant in the cytosol (Fig. 3 *D* and Fig. 4 *A*), neuronal autophagy activation most likely leads to AV recruitment of non-mitochondrial TBC1D15 in the cytosol through such a mechanism. As a result, mitochondrial bioenergetic stress-induced degradation of the cytosolic pool of TBC1D15 can disrupt TBC1D15 availability to AVs.

Several studies have highlighted the critical role of TBC1D15 in maintaining cellular homeostasis (73, 104–109). For instance, cardiac-specific deletion of *TBC1D15* in mice augmented phenotypes associated with myocardial ischemia/reperfusion (I/R) injury (109). Besides, TBC1D15 OE has shown protective effects *in vitro* in Parkinson’s disease patient-derived *GBA1*-linked neurons (8) and cardioprotection against myocardial I/R injury (108, 109). However, the role of TBC1D15 in AD and other tauopathies is poorly understood. The current study has demonstrated TBC1D15 deficiency in tauopathy brains. More importantly, normalization of TBC1D15 levels attenuates tau pathology, provides neuroprotection, and mitigates learning and memory deficits in tauopathy mice. Therefore, the current study fills an important knowledge gap regarding a crucial role of TBC1D15 in tauopathy-related pathologies. Given that mitochondrial/metabolic disruptions and autophagy defects are also prominent features in the amyloid pathology of AD (31, 96, 110–116), in the future, it is important to directly address whether AV-Mito contact dynamics is altered and contributes to autophagy dysfunction in these AD neurons.

Besides AV-Mito contact release defects, other mechanisms may also underlie TBC1D15 deficiency-induced autophagy defects in tauopathy and may work in an additive or synergistic manner. For example, studies from several groups indicate that TBC1D15 regulates lysosomal activity in non-neuronal cells (9, 73, 103, 105). Indeed, alterations in lysosomes were reported in tauopathy (28, 117–119). Autophagy is a lysosome-dependent pathway, and mature lysosomes with full activity of lysosomal hydrolases are mainly located in the soma of neurons (96, 97, 116). Hence, TBC1D15 deficits may also disrupt autophagy activity by compromising lysosomal function in the soma of tauopathy neurons. Additionally, previous studies have suggested that TBC1D15 plays a role in regulating mitophagy for mitochondrial maintenance in non-neuronal cells (9, 73, 103, 105). In fact, defective mitophagy has also been implicated in tauopathy (118, 119). Thus, TBC1D15 deficiency could contribute to mitochondrial defects by impairing mitophagy-mediated quality control of mitochondria. These potential TBC1D15 deficiency-related mechanisms may also underlie tauopathy-related autophagy/mitophagy deficits and need to be addressed further in the future.

Neuronal autophagy has been extensively studied, and an enormous body of work has unraveled the mechanisms underlying compartmentalized AV formation and maturation, the molecular machinery involved, the pathways governing cargo selection, degradation, and AV retrograde transport (46, 48, 49, 120–124). We have demonstrated that AV-Mito contacts impact AV retrograde movement and subsequent autophagic cargo clearance. While AV-Mito hyper-tethering can sufficiently accumulate autophagic cargos in normal axons, correction of such defects results in enhanced autophagic clearance and reduced tau burden in tauopathy axons and mouse brains (Fig. 4 *I-L*, *Supplemental* Fig. S6 *G-J,* and Fig. 5 *C-H*). Hence, the current study uncovers mitochondria-AV interactions as a new mechanism regulating neuronal autophagy by modulating AV retrograde transport. In addition, our recent studies have demonstrated that metabolically active mitochondria can supply membranes for AV biogenesis in neurons (42). Given that excessive AV-Mito tethering occurs in bioenergetically stressed mitochondria, this reduces the likelihood that the observed hyper-tethering is caused by mitochondria-enhanced AV biogenesis. Moreover, AVs in prolonged contact with mitochondria are amphisomes in nature after fusion with late endosomes (Fig. 2 *A* and *B*), indicating that AV tethering to mitochondria is unlikely due to incomplete AV formation is unlikely. A growing body of evidence suggests that mitochondria-organelle interactions play a crucial role in regulating mitochondrial dynamics and morphology, Ca^2+^ signaling, and the exchange of lipids and metabolites (3, 5). Thus, the physiological roles of AV-Mito contacts and how these roles are regulated under mitochondrial stress also require further investigation.

In normal neurons, AV-Mito contacts likely facilitate the exchange of ions and lipids. Recent studies have shown that AV/amphisome-localized transient receptor potential cation channel mucolipin subfamily member 1 (TRPML1) channels mediate Ca^2+^ efflux, enabling AVs/amphisomes to participate in local Ca^2+^ signaling (125). It is known that a proper Ca^2+^ load is crucial for mitochondrial function, including oxidative phosphorylation (OXPHOS) activity (126). AV-Mito contact sites may serve as a signaling platform allowing the modulation of mitochondrial Ca^2+^ homeostasis and energy metabolism. In addition, given that autophagy plays a major role in lipid breakdown, known as lipophagy, AV-Mito contacts may facilitate the AV-to-mitochondria transfer of lipids, such as fatty acids, thereby driving mitochondrial ATP production via β-oxidation. Indeed, emerging evidence indicates that fatty acids are an important fuel for synaptic function in the brain (127, 128). Therefore, AV-Mito contacts could act as a critical regulator of synaptic activity by supporting neuronal bioenergetics. Besides, while synaptic activation has been shown to enhance autophagy by promoting AV biogenesis (129, 130), synaptic activity also triggers AMPK activation (63, 131). Thus, it would be intriguing to assess the impact of synaptic activity on AV-Mito contact dynamics and vice versa. Taken together, ion and lipid transfer could constitute plausible physiological functions of AV-Mito contacts, which are likely involved in regulating mitochondrial activity and neuronal function.

In summary, the present study delineates a new class of inter-mitochondrial interactions through AV-Mito contacts that exhibit untethering defects in tauopathy neurons. Our work also advances current knowledge by providing new mechanistic insights into the role of AV-Mito contact dysregulation in autophagy dysfunction in the context of tauopathy. These findings have implications for understanding not only tauopathy diseases, but also other neurodegenerative disorders and aging-related phenotypes associated with mitochondrial defects, autophagy dysfunction, and toxic accumulation. The results also suggest TBC1D15-regulated AV-Mito contact-modulated autophagy activity as a promising therapeutic target for AD and other tauopathies.

## Materials and Methods

### Animals and Animal Care

TauP301S (PS19) and tauP301L (rTg4510) mouse lines (18, 132) were purchased from The Jackson Laboratory. All animal procedures were carried out in accordance with NIH guidelines and were approved by the Rutgers Institutional Animal Care and Use Committee (IACUC). The animal facilities at Rutgers University are fully accredited by the Association for Assessment and Accreditation of Laboratory Animal Care (AAALAC).

### Human brain specimens

Postmortem brain specimens from FTD patients and age-matched control subjects were obtained from the Harvard Tissue Resource Center and the Human Brain and Spinal Fluid Resource Center at UCLA. The specimens were from the frontal cortex and were quick-frozen (BA9). Seven control subjects and eight patient brains with postmortem intervals of 6.38 hr – 19.55 hr were used for analysis.

### Materials

Sources of antibodies or reagents are as follows: polyclonal antibodies against LC3 (Cell Signaling Technology, Cat# 2775), AMPKα (Cell Signaling Technology, Cat# 2532), p62/SQSTM1 (Abnova, Cat# H00008878-M01), NeuN (Millipore/Sigma, Cat# ABN78), TBC1D15 (Abcam, Cat# ab121396), Tau5 (DAKO, Cat# A0024), and synaptophysin/SYP (Abcam, Cat# ab32127); monoclonal antibodies against phospho-AMPKα (Cell Signaling Technology, Cat# 2535), AT8 (ThermoFisher Scientific, Cat# MN1020), syntaxin 1/STX1 (Santa Cruz Biotechnology, Cat# sc-12736), Rab7 (Sigma, Cat# R8779), GAPDH (Sigma, Cat# CB1001), TOM20 (Abcam, Cat# ab186734), mCherry (Takara, Cat# 632543), and Alexa fluor 488- (Cat# A-11017; Cat# A-11070), 546- (Cat# A-11018; Cat# A-11071), and 647- (Cat# A-21237; Cat# A-21246) conjugated secondary antibodies (Invitrogen); 10-NCP (VWR, Cat# 80017-188), trehalose (Cat# T0167), cycloheximide (Cat# 01810), rotenone (Cat# 557368), antimycin A (Cat# 1397-94-0), AICAR (Cat# A9978), CC (Cat# 171264), epoxomicin (Cat# 324800), CID 1067700 (Cat# SML0545), DMSO (Cat# D2650), and Sarkosyl (Cat# L9150) (Sigma); TBC1D15 shRNA (Cat# sc-154093-SH), Fis1 shRNA (Cat# sc-60644-SH), and control shRNA (Cat# sc-108060) (Santa Cruz Biotechnology); MitoView^TM^ Green (Cat# 70054) (biotium); LentiBrite RFP-LC3 Lentiviral Biosensor (Cat# 17-10143) (Sigma). The constructs encoding GFP-LC3, DsRed-Mito, and mRFP-LC3 were prepared as we previously described (41, 51, 75, 133). mCherry-TBC1D15 and mCherry-TBC1D15 D397A were from Vector Biolabs. Mito-ABKAR, Tom20-mChF-AIP, Tom20-mChF-AIP(TA), LAMP-mChF-AIP, and AIP(WT)-mChF-giantin were from Takanari Inoue, pEF6-myc-TBC1D15 was from Aimee Edinger, pEBFP2-Mito-7 was from Michael Davidson, mRFP-Rab7 was from Ari Helenius, EGFP-Rab7A, EGFP-Rab7A T22N, and EGFP-Rab7A Q67L were from Qing Zhong, EGFP-TBC1D15 was from Ivan Dikic.

### Primary neuronal cultures

Mouse cortical neurons were prepared from cortical tissues dissected from E18-19 mouse embryos (sex: random) using the Papain method as previously described (51, 78, 134–137). In brief, after dissociation by papain (Worthington), cortical neurons were resuspended in a plating medium (5% FBS, insulin, glutamate, G5 and 1 × B27) supplemented with 100 × L-glutamine in Neurobasal medium (Invitrogen) and plated at a density of 100,000 cells per cm^2^ on polyornithine- and fibronectin-coated coverslips (Carolina Cover Glasses) or 35 mm dishes. Twenty-four hours after plating, neurons were maintained in conditioned medium with half-feed changes of neuronal feeding medium (1 × B27 in Neurobasal medium) every 3 days. Primary cortical neurons were cultured from breeding mice of tauP301S Tg line with non-Tg controls (132). Genotyping assays were performed following culture plating to verify mouse genotypes. We examined both tauP301S Tg and non-Tg neurons derived from their littermates. Non-Tg and tauP301S Tg neurons were transfected with various constructs at DIV5-7 using Lipofectamine 2000 (Invitrogen), followed by time-lapse imaging 5-10 days after transfection prior to quantification analysis. For biochemical analysis, harvested neurons were lysed with ice-cold lysis buffer [1% Triton X-100, 8 μg/ml aprotinin, 10 μg/ml leupeptin, 0.4 mM PMSF in TBS with 1 × PhosSTOP (Millipore/Sigma, Cat# 4906837001)] and then proceeded to centrifugation and protein quantification.

### Determination of mitochondria-localized AMPK signaling in live neurons

Mito-ABKAR was transfected into cortical neurons at DIV5-6 using Lipofectamine 2000. The neurons were imaged using an Olympus FV3000 microscope at DIV13-14. The FRET signal (YFP/CFP emission ratio) was measured and calculated from individual cells expressing Mito-ABKAR, reflecting mitochondrial AMPK activity.

### Cell fractionation

The cytosol and mitochondria-enriched fractions were prepared as previously described (75, 133, 135, 137). Briefly, cortical neurons were cultured in 100-mm plates at a density of 5 × 10^6^ in the presence and absence of glutamine for 24 h at DIV14. After washing once with PBS (Quality Biological, 114-058-131), cells were harvested and suspended in ice-cold Isolation Buffer (IB) [10 mM Tris-HCl, 1 mM EGTA, 1 mM EDTA, 0.25 M sucrose, and protease inhibitors (Roche, 4693159001), pH7.4]. Cells were then homogenized by passing through a 25-gauge needle 20 times using a 1-ml syringe on ice. Following centrifugation at 1,000 × g for 10 min at 4°C, the supernatant was saved as post-nuclear supernatant (PNS). PNS was centrifuged at 15,000 × g for 10 min to separate the mitochondria-enriched fraction (Mito) from the cytosol-enriched fraction (Sup). The same amount of protein (10 μg) from each fraction was resolved by 4-12% Bis-Tris PAGE for sequential western blots on the same membranes after stripping between each application of antibody.

### Brain tissue extraction

Human or mouse brain tissues were lysed with ice-cold lysis buffer [1% Triton X-100, 8 μg/ml aprotinin, 10 μg/ml leupeptin, 0.4 mM PMSF in TBS with phosphatase inhibitors (1 × PhosSTOP) (Millipore/Sigma, Cat# 4906837001)]. For obtaining sarkosyl-insoluble tau, tauP301S Tg mouse brain tissues were homogenized in 10 volumes of high salt/sucrose extraction buffer [25 mM Tris/HCl pH7.4, 0.8 M NaCl, 10% sucrose, 1mM EGTA, a protease inhibitor cocktail (Sigma, Cat# 11836170001), and 1 × PhosSTOP]. The homogenates were centrifuged at 20,000 × g for 20 min at 4°C to obtain supernatant (S1) and pellet (P1) fractions. Pellets were homogenized in 5 volumes of high salt/sucrose buffer and centrifuged as above. The resulting pellets were discarded, and the supernatants (S2) were incubated with sarkosyl (1% final concentration; Sigma) for 1 h at 37°C, followed by centrifugation in a Beckman TLA100.2 rotor at 150,000 × g for 1 h at 4 °C to obtain salt and sarkosyl-extractable (S3) and sarkosyl-insoluble (P3) fractions. The supernatants (S3) were collected, while the P3 pellet was re-suspended in TBS containing protease and phosphatase inhibitors [25 mM Tris/HCl (pH 7.4), 137 mM NaCl, a protease inhibitor cocktail, and 1 × PhosSTOP, followed by protein quantification by BCA assay.

### Western blot

Protein quantification of cortical neuron lysates or tissue homogenates was performed by BCA assay (Pierce Chemical Co.). Equal amounts of proteins were loaded and resolved by 4-12% Bis-Tris NuPAGE protein gel and analyzed by sequential western blots on the same membranes after stripping between each application of antibody. For semi-quantitative analysis, protein bands detected by ECL were scanned into Adobe Photoshop 2025 and analyzed using NIH ImageJ. Care was taken during exposure of the ECL film to ensure that the intensity readouts were within the linear range of the standard curve blot detected by the same antibody.

### Live neuron imaging

Neurons were transferred to Tyrode’s solution containing 10 mM HEPES, 10 mM glucose, 1.2 mM CaCl_2_, 1.2 mM MgCl_2_, 3 mM KCl and 145 mM NaCl, pH 7.4. The temperature was maintained at 37°C with an air stream incubator. Cells were visualized with a 60 × oil immersion lens (1.3 numerical aperture) on an Olympus FV3000 confocal microscope, using 405 nm excitation for CFP or BFP, 488 nm for GFP, and 559 nm for DsRed, mRFP, or mCherry. We selected the axons with at least 150 μm in length and more than 200 μm away from the soma of non-Tg or tauP301S Tg neurons. Axonal processes were selected as we previously reported (41, 75, 134–136). Briefly, axons in live images were distinguished from dendrites based on known morphologic characteristics: greater length, thin and uniform diameter, and sparse branching (138). Only those that appeared to be single axons and separate from other processes in the field were chosen for recording axonal mitochondrial transport. Regions where crossing or fasciculation occurred were excluded from the analysis.

For the analysis of GFP-p62 puncta/clusters, a minimum threshold intensity determined by the thresholding function of NIH ImageJ was subtracted from each image prior to measurement. The number of p62 puncta/clusters with a size ranging from 0.15 μm^2^ to 0.50 μm^2^ was then counted within the axonal processes in the background-subtracted images.

For time-lapse imaging, time-lapse sequences of 1,024 × 1,024 pixels (8 bit) were collected at 3-sec intervals with 1% intensity of the laser to minimize laser-induced bleaching and cell damage while maximizing pinhole opening. Time-lapse images were captured for a total of 100 frames for single-channel imaging. For dual- or triple-channel imaging to examine AV-Mito contact dynamics, 300 frames were collected. Recordings were started 6 minutes after the coverslip was placed in the chamber. The stacks of representative images were imported into NIH ImageJ software and converted to kymographs. To trace the anterograde or retrograde movement of axonal AVs and to count stationary ones, kymographs were generated as described previously (135, 139) with extra plug-ins for NIH ImageJ. Briefly, we used the “Straighten” plugin to straighten curved axons and the “Grouped ZProjector” to z-axially project re-sliced time-lapse images. The height of the kymographs represents recording time, while the width represents the length (μm) of the axon imaged. Counts were averaged from 100 frames for single-channel time-lapse images and from 300 frames for imaging AV-Mito contacts to ensure the accuracy of stationary and motile events. Vertical lines represent stationary organelles, while slanted lines to the right (negative slope) represent anterograde movement, and to the left (positive slope) indicate retrograde movement. An AV was considered stationary if its displacement was less than 5 μm. Relative motility of vesicles or organelles is described as the percentage of anterograde, retrograde, or stationary events of total vesicles or organelles.

### Grid preparation for primary tauP301S Tg cortical neurons

Quantifoil R1/4 200-mesh SiO_2_ gold Finder, R2/2 200-mesh extra thick carbon gold NH_2_ Finder, and R2/2 300-mesh SiO_2_ gold cryo-EM grids (Quantifoil Micro Tools GmbH) were glow-discharged to render surface hydrophilicity and briefly dipped into 70% ethanol to sterilize. Prior to cell plating, these grids were coated with polyornithine and fibronectin and washed three times with PBS. The grids were then plated with isolated cortical neurons derived from tauP301S Tg mouse brains as described above. Neurons were maintained in conditioned medium with half-feed changes of neuronal feeding medium (1 × B27 in Neurobasal medium) every 3 days and treated at DIV13 with trehalose overnight before vitrification. To label AVs and mitochondria, neurons were transduced with LentiBrite RFP-LC3 Lentiviral Biosensor (Sigma) and loaded with MitoView^TM^ Green (biotium). TauP301S Tg neurons grown on grids were vitrified using a Leica EM GP plunger (Leica Microsystems) operating at 95% humidity at 20°C. Grids were back-blotted for 10 seconds before plunging them into liquid ethane. The vitrified grids were transferred into grid storage boxes and stored in clean liquid nitrogen.

### Correlative light and electron microscopy (CLEM) and cryo-electron tomography (cryo-ET) data collection

Grids were loaded into a Leica DM6 FS microscope equipped with a 50 × objective and a DFC 365 FX camera. The microscope stage was kept at −195°C with liquid N_2_ vapor. Fluorescence and corresponding bright field montage images of the grids were acquired using the Leica LASX software with a GFP filter set (excitation 470/40 nm, dichroic 495 nm, emission 525/50 nm) and a TXR filter set (excitation 560/40 nm, dichroic 585 nm, emission 630/76 nm). Areas of the grid with colocalization of MitoView Green and mRFP densities were marked as targets for cryo-ET imaging.

Grids carrying tauP301S Tg neurons were then loaded into a 300 kV Titan Krios G3i TEM equipped with a K3 detector (Gatan, Inc.) and a BioQuantum energy filter set to 20 eV slit width. A low-magnification montage of the entire grid was first collected and correlated with cryo-FLM images of the same grid to identify areas containing axonal processes with potential AV-Mito contacts for tilt series data collection (Tomography version 5.23, Thermo Scientific). Target areas were images at 3,600 × magnification to identify candidate AV-Mito contact sites. Tilt series of the selected areas were collected at 19,500 × magnification (pixel size 4.59 Å/pixel), or 26,000 × magnification (pixel size 3.42 Å/pixel). The imaging settings used were spot size 8, a 50 μm condenser aperture, a 100 μm objective aperture, and approximately −8 μm defocus. Tilt series were collected from -60° to 60° with 3° increments using a continuous schema in counting mode, with a total dose of ∼120–160 e/Å2 per tilt series.

### Cryo-ET data processing

Tilt series alignment and reconstruction were performed using IMOD (140). Membranes were automatically segmented using the TomoSegMemTV package (141). Manual refinement of the segmentation results and visualization of cellular features were performed using UCSF Chimera (University of California, San Francisco) (142).

### Tissue preparation and immunohistochemistry

Animals were anesthetized with 2.5% avertin (0.35ml per mouse) and transcardially perfused with fixation buffer (4% paraformaldehyde in PBS, pH 7.4). Brains were dissected out and postfixed in fixation buffer overnight and then placed in 30% sucrose at 4°C. Ten-µm-thick coronal sections were collected consecutively to the level of the hippocampus and used to study the co-localization of various markers. After incubation with blocking buffer (5% goat serum, 0.3% Triton X-100, 3% BSA, 1% glycine in PBS) at room temperature (RT) for 1 hr, the sections were incubated with primary antibodies at 4°C overnight, followed by incubation with fluorescent secondary antibodies at 1:600 dilution at RT for 1 hr. After fluorescence immunolabeling, the sections were stained with DAPI and washed three times in PBS. The sections were then mounted with an anti-fading medium (vector laboratories, Cat# H-5000) for imaging. Confocal images were obtained using an Olympus FV3000 oil-immersion 40× objective with a sequential-acquisition setting. Eight to ten sections were taken from the top to the bottom of the specimen, and the brightest point projections were made. Fluorescence intensities of AT8, PHF1, synaptophysin (SYP), NeuN, or p62/SQSTM1 were expressed in arbitrary units of fluorescence per square area. The mean intensities of AT8, PHF1, SYP, NeuN, or p62/SQSTM1 in the hippocampi of tauP301S Tg mouse brains were normalized to those in the brains of control non-Tg littermates or tauP301S Tg mice injected with AAV-mCherry. Data were obtained from at least three pairs of tauP301S Tg mice, and the number of imaging brain slices used for quantification is indicated in the figures.

### Transmission electron microscopy

Hippocampi from 8-month-old non-Tg and tauP301S Tg mouse brains were cut into small specimens (one dimension < 1 mm) and fixed in Trump’s fixative (Electron Microscopy Sciences) for 2 hr at RT. The sections were then washed with 0.1 M Cacodylate buffer and postfixed in 1% osmium tetroxide, followed by dehydrating in ethanol and embedding using the EM bed 812 kit (Electron Microscopy Sciences) according to a standard procedure. Images were acquired using an electron microscope (1200EX; JEOL) (Electron Imaging Facility, Department of Pathology and Laboratory Medicine, Robert Wood Johnson Medical School). For quantitative studies, the percentage of autophagic vacuole (AV)-like organelles and the number of AVs or mitochondria in AV-Mito contacts at presynaptic terminals were counted from electron micrographs, respectively. AV-like structures were characterized by double-membrane structures containing other organelles, vesicles, or cytoplasmic material (134, 143, 144). The analyses of AV-Mito contacts were based on the guidelines for defining membrane contact sites (145). Specifically, AV-Mito contact sites were defined as sites of contact within a reciprocal distance of 80 nm (*Supplemental* Fig. S1 *A*). The corresponding intermembrane distances and lengths between the two organelles at individual AV-Mito contact sites, as well as their frequencies, were quantified and compared with those in control groups. Quantification analysis was performed blindly to the condition.

### Real time qRT-PCR measurements

Total RNAs were extracted from mouse or human brain tissues using Trizol (Invitrogen Co., Carlsbad, CA) according to the manufacturer’s instructions. PCR was performed in an Eco real-time qPCR system (Illumina, San Diego,CA) using iScriptTM One-Step RT-PCR Kit With SYBR Green (Bio-Rad Laboratories Inc., Hercules, CA) in 20 µL reactions with 20 ng of RNA template. We used forward (5′- GTC TCC TCT GAC TTC AAC AGC G -3′) and reverse (5′- ACC ACC CTG TTG CTG TAG CCA A -3′) primers for human GAPDH, and forward (5′-ACC ACA GTC CAT GCC ATC AC -3′) and reverse (5′- TCC ACC ACC CTG TTG CTG TA -3′) primers for mouse GAPDH, respectively. We used forward (5′- CCA CAT ACC AAT GGA GAT GCT CC -3′) and reverse (5′- ATA GGA CCA GCC CAT ACC CTC T -3′) primers for human TBC1D15, and forward (5′- CAG GAG AAG AGA AAC TCA CGC C -3′) and reverse (5′- TGT CGT GAA GCA TCA GCC C -3′) primers for mouse TBC1D15, respectively. Samples were measured two to three times and average values were used for the calculation of relative fold changes. The relative levels of TBC1D15 mRNA were normalized to GAPDH mRNA levels in each preparation. For each experiment, the values for non-Tg or human control subjects were set to 1, and other values were normalized accordingly.

### AAV Design and injection

The AAV constructs were built using standard molecular biology techniques. All constructs were self-complementary adeno-associated virus (AAV) genomes with a Chicken Beta Actin hybrid promoter and bovine growth hormone poly-adenylation signal. The upstream cDNA in both constructs was mCherry with a mutated stop codon. In the control construct, a stop codon was engineered 3’ to the 2A sequence and no second gene was inserted. The TBC1D15 or TBC1D15 D397A mutant construct contained the human cDNA in frame 3’ to the 2A sequence. The AAV2/9-mCherry, AAV2/9-mCherry-hTBC1D15, and AAV2/9-mCherry-hTBC1D15 D397A mutant viruses were produced by Vector BioLabs. Adult non-Tg and tauP301S Tg mice at 2 months of age were injected into the hippocampus (AP: − 2 mm, LAT: − 1.5 mm, DV: + 1.75mm) and the cortex (AP: + 1.5 mm, LAT: + 1.5 mm, DV: + 1.5 mm) of both cerebral hemispheres according to the stereotaxic atlas of Franklin and Paxinos (2001) using 4-8 × 10^9^ total viral particles per side and analyzed 4-8 months after injection (*Supplemental* Fig. S6 *A* and Fig. S7).

### Behavioral assays

Only male mice were assessed for cognitive and emotional behavior. Groups contained 10-17 male mice/group for each behavioral task. Tests were performed in mice aged 6-7 months. All tests used the ANY-maze Video Tracking System (Stoelting Co.) with additional video review by experimenters who were blind to treatment group conditions.

#### Sociability and preference for social novelty test

The test for sociability and preference for social novelty was conducted as previously described (87–90). The apparatus consisted of a rectangular, three-chambered box. Each chamber was measured 26.5 × 46 × 30 cm with the dividing walls made of white Plexiglas and with small square openings (4 × 4 cm) allowing access into each chamber. The stranger mouse (stranger 1 (S1), a 7-month-old male C57BL/6 mouse that was not a littermate nor previously used in any other experiments) was enclosed in a small, circular metal wire cage (10 × 10 cm) that allowed nose contact between the bars but prevented fighting. For the first 10-min session, the placement of S1 in the left or right-side chamber was systematically alternated between trials, with the opposite side chamber left empty (E). The test mouse was first placed in the middle chamber and allowed to explore the entire social test box. Using the ANY-maze Video Tracking System, the movement of the test mouse and the amount of time that the test mouse spent in each chamber were tracked for 10 min for subsequent data analysis to quantify social preference for S1. After the sociability test session was over, there was a 10-min inter-trial interval before the test mouse was guided to the center chamber for a social recognition test in a second 10-min session. Also, prior to this session, a second, unfamiliar male mouse (stranger 2 (S2), 7 months old and not a littermate) was introduced into the empty side chamber. This S2 mouse was enclosed in a small wire cage identical to that for the S1 mouse. The test mouse had a choice between the first, already-investigated familiar mouse (S1) and the novel unfamiliar mouse (S2). As described above, the amount of time spent in each chamber with S1 or S2 during this second 10-min session was recorded for the measurement of social recognition memory. Thus, sociability and recognition memory were assessed in trials by analyzing the amount of time spent exploring the empty chamber E or the chamber with S1 in the first session and the chambers containing S1 or S2 in the second session, respectively.

#### Morris water maze (MWM)

The water maze consisted of a pool (110 cm in diameter) containing tap water (24°C ± 1°C) made opaque with non-toxic powdered paint. A platform (9.4 cm in diameter) was submerged 1.5 cm beneath the water surface. The pool was surrounded by a curtain and a facing wall. Distinct cues were placed on the curtain and wall and remained the same for all animals. Mice were trained to locate the hidden platform in a single day (92), with a probe memory trial given after the final acquisition trial. The training was conducted in four blocks. The first block (Block 1) of trials consisted of three trials in which the target platform was made visible by an affixed flag. These three flagged trials ensured that animals were primed to seek an escape platform, as well as tested for any visual deficits. Blocks 2-4 consisted of hidden-platform training, with each block consisting of four trials, with the platform hidden below the surface, off-center and randomly located (across animals) in one of the four quadrants of the circular pool. For each animal, the location of the platform was held constant across trials but was randomized across animals within each treatment group. Each platform location was equally represented across all treatment groups. For each training block, the inter-trial interval was 5 min, such that each mouse completed a block in approximately 20 minutes. For each trial in a block, mice were placed into the water with the head facing the tank wall and allowed a maximum of 60 sec to find the hidden platform (mice were returned to their holding cage until the next trial if they had not found the platform by 60 seconds). Each succeeding trial within a block involved a different starting location (north, south, west, or east relative to the platform). Once a block was completed, it was repeated for each mouse, so that all mice were run through three hidden-platform training blocks (Blocks 2-4), constituting a total of 12 acquisition trials. One hour or 24 hours after the last trial of Block 4, a 60-second probe trial was conducted to test for platform location memory. In the probe trial, the platform was not in the pool, with the 60-second period being used to measure the total time spent swimming in the target quadrant (usual platform location) relative to the remaining quadrants. Data consisted of the time to locate the hidden platform during learning, the time spent in the target quadrant during the probe test, and the path length during the probe test. These data were calculated by the ANY-maze Video Tracking System.

#### Contextual fear conditioning

Conditioning was conducted in Coulbourn operant learning chambers that measured 19 × 20.3 × 30.5 cm and were located inside sound-attenuating cabinets (Coulbourn Instruments, Whitehall, PA). The chamber contained a steel grid floor connected to a programmable shocker, while the walls were made of clear Plexiglas. For contextual fear conditioning, the mice were placed within the conditioning chamber for 3 min to develop a representation of the context prior to the onset of a single unconditioned stimulus (US; 1.0 mA footshock; 1 second duration). Following the shock, they were allowed to remain in the chamber for 2 min, during which immediate freezing was measured continuously. Mice were then returned to the home cages. Memory was tested 24 hours after training for 4 min in the same conditioning chamber. Animal movements were tracked with the ANY-maze Video Tracking System.

### Statistical analysis

Statistical parameters, including the definitions and exact value of n (e.g., number of neurons, number of axons, number of AV-Mito contacts, number of AVs or mitochondria, number of mouse brain slice sections, number of animals, etc), p values, and the types of the statistical tests were reported in the figures and in the corresponding figure legends. For normalization, in studies with non-Tg and tauP301S Tg groups, the data points from the tauP301S Tg group were normalized to those in the non-Tg group. For experiments using non-Tg or tauP301S Tg alone, the data were normalized to a basal condition, a vehicle-treated condition, or a control plasmid condition. For studies with AAV-injected mice, the data points were normalized to those in non-Tg mice or tauP301S Tg mice injected with AAV-mCherry. Additionally, for experiments using the brains of tauP301L Tg mice or FTD patients, the data were normalized to those in control mice or control human subjects. In all biochemical studies, data points were first normalized to loading controls from the same sample and then to the control condition within the same experiment. Statistical analysis was carried out using GraphPad Prism (version 10.6) or IBM SPSS (version 31.0). All statistical comparisons were conducted on data originating from three or more biologically independent experimental replicates. Statistical comparisons between two groups were performed by an unpaired student’s *t*-test. Comparisons between three or more groups were performed using one-way or two-way ANOVA with post hoc testing with Tukey’s or Sidak’s correction unless otherwise indicated. For comparison of group means between treatments or other experimental conditions with multiple observations made from a shared source (e.g., a culture or a brain slice), linear mixed-effects (LME) analysis was conducted (146). Specifically, while the treatment or condition of interest was set as a fixed effect, the shared source was modeled as a random effect to account for data dependence, thus controlling for type I error. For statistically significant fixed effects involving more than 2 groups, follow-up analyses using the Bonferroni correction were performed for group mean comparisons. The LME analysis was carried out using IBM SPSS (version 31.0). Data are expressed as mean ± SEM. Differences were considered significant with *P* < 0.05.

## Data and materials availability

Tomograms of AV-Mito contacts in tauP301S Tg cortical neurons have been deposited at the EMDB repository under accession numbers EMD-75946, EMD-75947, and EMD-75949, and are publicly available as of the date of publication.

## Acknowledgments

We are grateful to Elaine Gavin, You Cai, and Selina Wadhera for their research assistance and constructive discussion; Rajesh Patel at the EM facility in the Department of Pathology and Laboratory Medicine, Robert Wood Johnson Medical School, and Jason Kaelber and Ashley Bernstein at the Rutgers CryoEM & Nanoimaging Facility for technical help; Takanari Inoue for Mito-ABKAR, Tom20-mChF-AIP, Tom20-mChF-AIP(TA), LAMP-mChF-AIP, and AIP(WT)-mChF-giantin plasmids, Aimee Edinger for pEF6-myc-TBC1D15 plasmid, Michael Davidson for pEBFP2-Mito-7 plasmid, Ari Helenius for mRFP-Rab7, Qing Zhong for EGFP-Rab7A, EGFP-Rab7A T22N, and EGFP-Rab7A Q67L plasmids, Ivan Dikic for EGFP-TBC1D15 plasmid, and Noboru Mizushima for GFP-p62 plasmid; Peter Davies for PHF1 antibody. This research was supported by National Institutes of Health grants R01NS089737, RF1NS130881, R21AG089974 (to Q.C.), and National Science Foundation CAREER MCB-2046180 (W.D.).

## References

1. T. F. Gendron, L. Petrucelli, The role of tau in neurodegeneration. Mol Neurodegener 4, 13 (2009).

2. E. M. Mandelkow, E. Mandelkow, Biochemistry and cell biology of tau protein in neurofibrillary degeneration. Cold Spring Harb Perspect Med 2, a006247 (2012).

3. Y. C. Wong, S. Kim, W. Peng, D. Krainc, Regulation and Function of Mitochondria-Lysosome Membrane Contact Sites in Cellular Homeostasis. Trends Cell Biol 29, 500–513 (2019).

4. S. Vrijsen et al., Inter-organellar Communication in Parkinson’s and Alzheimer’s Disease: Looking Beyond Endoplasmic Reticulum-Mitochondria Contact Sites. Front Neurosci 16, 900338 (2022).

5. M. Petkovic, C. E. O’Brien, Y. N. Jan, Interorganelle communication, aging, and neurodegeneration. Genes Dev 35, 449–469 (2021).

6. J. Cisneros, T. B. Belton, G. C. Shum, C. G. Molakal, Y. C. Wong, Mitochondria-lysosome contact site dynamics and misregulation in neurodegenerative diseases. Trends Neurosci 45, 312–322 (2022).

7. S. Kim, R. Coukos, F. Gao, D. Krainc, Dysregulation of organelle membrane contact sites in neurological diseases. Neuron 110, 2386–2408 (2022).

8. S. Kim, Y. C. Wong, F. Gao, D. Krainc, Dysregulation of mitochondria-lysosome contacts by GBA1 dysfunction in dopaminergic neuronal models of Parkinson’s disease. Nat Commun 12, 1807 (2021).

9. Y. C. Wong, W. Peng, D. Krainc, Lysosomal Regulation of Inter-mitochondrial Contact Fate and Motility in Charcot-Marie-Tooth Type 2. Dev Cell 50, 339–354 e334 (2019).

10. Y. C. Wong et al., Misregulation of mitochondria-lysosome contact dynamics in Charcot-Marie-Tooth Type 2B disease Rab7 mutant sensory peripheral neurons. Proc Natl Acad Sci U S A 120, e2313010120 (2023).

11. E. Area-Gomez et al., A key role for MAM in mediating mitochondrial dysfunction in Alzheimer disease. Cell Death Dis 9, 335 (2018).

12. E. Area-Gomez et al., Presenilins are enriched in endoplasmic reticulum membranes associated with mitochondria. Am J Pathol 175, 1810–1816 (2009).

13. E. Area-Gomez et al., Upregulated function of mitochondria-associated ER membranes in Alzheimer disease. Embo j 31, 4106–4123 (2012).

14. T. Hamano et al., Autophagic-lysosomal perturbation enhances tau aggregation in transfectants with induced wild-type tau expression. Eur J Neurosci 27, 1119–1130 (2008).

15. U. Kruger, Y. Wang, S. Kumar, E. M. Mandelkow, Autophagic degradation of tau in primary neurons and its enhancement by trehalose. Neurobiol Aging 33, 2291–2305 (2012).

16. J. A. Rodriguez-Navarro et al., Trehalose ameliorates dopaminergic and tau pathology in parkin deleted/tau overexpressing mice through autophagy activation. Neurobiol Dis 39, 423–438 (2010).

17. Y. Wang et al., Tau fragmentation, aggregation and clearance: the dual role of lysosomal processing. Hum Mol Genet 18, 4153–4170 (2009).

18. K. Santacruz et al., Tau suppression in a neurodegenerative mouse model improves memory function. Science 309, 476–481 (2005).

19. A. B. Rocher et al., Structural and functional changes in tau mutant mice neurons are not linked to the presence of NFTs. Exp Neurol 223, 385–393 (2010).

20. J. L. Crimins, A. B. Rocher, J. I. Luebke, Electrophysiological changes precede morphological changes to frontal cortical pyramidal neurons in the rTg4510 mouse model of progressive tauopathy. Acta Neuropathol 124, 777–795 (2012).

21. M. Polydoro et al., Soluble pathological tau in the entorhinal cortex leads to presynaptic deficits in an early Alzheimer’s disease model. Acta Neuropathol 127, 257–270 (2014).

22. J. McInnes et al., Synaptogyrin-3 Mediates Presynaptic Dysfunction Induced by Tau. Neuron 97, 823–835 e828 (2018).

23. B. R. Hoover et al., Tau mislocalization to dendritic spines mediates synaptic dysfunction independently of neurodegeneration. Neuron 68, 1067–1081 (2010).

24. H. C. Tai et al., Frequent and symmetric deposition of misfolded tau oligomers within presynaptic and postsynaptic terminals in Alzheimer’s disease. Acta Neuropathol Commun 2, 146 (2014).

25. K. V. Kuchibhotla et al., Neurofibrillary tangle-bearing neurons are functionally integrated in cortical circuits in vivo. Proc Natl Acad Sci U S A 111, 510–514 (2014).

26. C. A. Lasagna-Reeves et al., Tau oligomers impair memory and induce synaptic and mitochondrial dysfunction in wild-type mice. Mol Neurodegener 6, 39 (2011).

27. N. Watamura et al., In vivo hyperphosphorylation of tau is associated with synaptic loss and behavioral abnormalities in the absence of tau seeds. Nat Neurosci 10.1038/s41593-024-01829-7 (2024).

28. A. Piras, L. Collin, F. Gruninger, C. Graff, A. Ronnback, Autophagic and lysosomal defects in human tauopathies: analysis of post-mortem brain from patients with familial Alzheimer disease, corticobasal degeneration and progressive supranuclear palsy. Acta Neuropathol Commun 4, 22 (2016).

29. B. Caballero et al., Interplay of pathogenic forms of human tau with different autophagic pathways. Aging Cell 17 (2018).

30. R. H. Swerdlow, J. M. Burns, S. M. Khan, The Alzheimer’s disease mitochondrial cascade hypothesis. J Alzheimers Dis 20 **Suppl 2**, S265–279 (2010).

31. G. E. Gibson, Q. Shi, A mitocentric view of Alzheimer’s disease suggests multi-faceted treatments. J Alzheimers Dis 20 **Suppl 2**, S591–607 (2010).

32. Q. Cai, Y. Y. Jeong, Mitophagy in Alzheimer’s Disease and Other Age-Related Neurodegenerative Diseases. Cells 9 (2020).

33. P. H. Reddy, M. F. Beal, Amyloid beta, mitochondrial dysfunction and synaptic damage: implications for cognitive decline in aging and Alzheimer’s disease. Trends Mol Med 14, 45–53 (2008).

34. Q. Cai, P. Tammineni, Alterations in Mitochondrial Quality Control in Alzheimer’s Disease. Front Cell Neurosci 10, 24 (2016).

35. D. M. A. Oliver, P. H. Reddy, Molecular Basis of Alzheimer’s Disease: Focus on Mitochondria. J Alzheimers Dis 72, S95–S116 (2019).

36. W. Wang, F. Zhao, X. Ma, G. Perry, X. Zhu, Mitochondria dysfunction in the pathogenesis of Alzheimer’s disease: recent advances. Mol Neurodegener 15, 30 (2020).

37. X. Zhu, G. Perry, M. A. Smith, X. Wang, Abnormal mitochondrial dynamics in the pathogenesis of Alzheimer’s disease. J Alzheimers Dis 33 **Suppl 1**, S253–262 (2013).

38. M. Dumont et al., Behavioral deficit, oxidative stress, and mitochondrial dysfunction precede tau pathology in P301S transgenic mice. FASEB J 25, 4063–4072 (2011).

39. I. Lopez-Gonzalez et al., Neuroinflammatory Gene Regulation, Mitochondrial Function, Oxidative Stress, and Brain Lipid Modifications With Disease Progression in Tau P301S Transgenic Mice as a Model of Frontotemporal Lobar Degeneration-Tau. J Neuropathol Exp Neurol 74, 975–999 (2015).

40. K. J. Kopeikina et al., Tau accumulation causes mitochondrial distribution deficits in neurons in a mouse model of tauopathy and in human Alzheimer’s disease brain. Am J Pathol 179, 2071–2082 (2011).

41. Y. Y. Jeong et al., Broad activation of the Parkin pathway induces synaptic mitochondrial deficits in early tauopathy. Brain 10.1093/brain/awab243 (2022).

42. N. Jia et al., Mitochondrial bioenergetics stimulates autophagy for pathological MAPT/Tau clearance in tauopathy neurons. Autophagy 21, 54–79 (2025).

43. Y. Y. Jeong, N. Jia, D. Ganesan, Q. Cai, Broad activation of the PRKN pathway triggers synaptic failure by disrupting synaptic mitochondrial supply in early tauopathy. Autophagy 10.1080/15548627.2022.2039987, 1-3 (2022).

44. M. Escobar-Khondiker et al., Annonacin, a natural mitochondrial complex I inhibitor, causes tau pathology in cultured neurons. J Neurosci 27, 7827–7837 (2007).

45. T. Terada et al., Mitochondrial complex I abnormalities is associated with tau and clinical symptoms in mild Alzheimer’s disease. Mol Neurodegener 16, 28 (2021).

46. S. Lee, Y. Sato, R. A. Nixon, Lysosomal proteolysis inhibition selectively disrupts axonal transport of degradative organelles and causes an Alzheimer’s-like axonal dystrophy. J Neurosci 31, 7817–7830 (2011).

47. S. Maday, E. L. Holzbaur, Autophagosome biogenesis in primary neurons follows an ordered and spatially regulated pathway. Dev Cell 30, 71–85 (2014).

48. S. Maday, E. L. Holzbaur, Compartment-Specific Regulation of Autophagy in Primary Neurons. J Neurosci 36, 5933–5945 (2016).

49. X. T. Cheng, B. Zhou, M. Y. Lin, Q. Cai, Z. H. Sheng, Axonal autophagosomes recruit dynein for retrograde transport through fusion with late endosomes. J Cell Biol 209, 377–386 (2015).

50. A. K. Stavoe, S. E. Hill, D. H. Hall, D. A. Colon-Ramos, KIF1A/UNC-104 Transports ATG-9 to Regulate Neurodevelopment and Autophagy at Synapses. Dev Cell 38, 171–185 (2016).

51. P. Tammineni, X. Ye, T. Feng, D. Aikal, Q. Cai, Impaired retrograde transport of axonal autophagosomes contributes to autophagic stress in Alzheimer’s disease neurons. Elife 6 (2017).

52. T. Feng, P. Tammineni, C. Agrawal, Y. Y. Jeong, Q. Cai, Autophagy-mediated Regulation of BACE1 Protein Trafficking and Degradation. J Biol Chem 292, 1679–1690 (2017).

53. A. S. Tsvetkov et al., A small-molecule scaffold induces autophagy in primary neurons and protects against toxicity in a Huntington disease model. Proc Natl Acad Sci U S A 107, 16982–16987 (2010).

54. F. Guerra, C. Bucci, Multiple Roles of the Small GTPase Rab7. Cells 5 (2016).

55. J. O. Agola et al., A competitive nucleotide binding inhibitor: in vitro characterization of Rab7 GTPase inhibition. ACS Chem Biol 7, 1095–1108 (2012).

56. A. Mukhopadhyay, K. Funato, P. D. Stahl, Rab7 regulates transport from early to late endocytic compartments in Xenopus oocytes. J Biol Chem 272, 13055–13059 (1997).

57. K. Onoue et al., Fis1 acts as a mitochondrial recruitment factor for TBC1D15 that is involved in regulation of mitochondrial morphology. J Cell Sci 126, 176–185 (2013).

58. Y. C. Wong, D. Ysselstein, D. Krainc, Mitochondria-lysosome contacts regulate mitochondrial fission via RAB7 GTP hydrolysis. Nature 554, 382–386 (2018).

59. S. Herzig, R. J. Shaw, AMPK: guardian of metabolism and mitochondrial homeostasis. Nat Rev Mol Cell Biol 19, 121–135 (2018).

60. D. G. Hardie, AMP-activated/SNF1 protein kinases: conserved guardians of cellular energy. Nat Rev Mol Cell Biol 8, 774–785 (2007).

61. V. Vingtdeux, P. Davies, D. W. Dickson, P. Marambaud, AMPK is abnormally activated in tangle- and pre-tangle-bearing neurons in Alzheimer’s disease and other tauopathies. Acta Neuropathol 121, 337–349 (2011).

62. J. M. Corton, J. G. Gillespie, S. A. Hawley, D. G. Hardie, 5-aminoimidazole-4-carboxamide ribonucleoside. A specific method for activating AMP-activated protein kinase in intact cells? Eur J Biochem 229, 558–565 (1995).

63. S. Li, G. J. Xiong, N. Huang, Z. H. Sheng, The cross-talk of energy sensing and mitochondrial anchoring sustains synaptic efficacy by maintaining presynaptic metabolism. Nat Metab 2, 1077–1095 (2020).

64. J. Kim, M. Kundu, B. Viollet, K. L. Guan, AMPK and mTOR regulate autophagy through direct phosphorylation of Ulk1. Nat Cell Biol 13, 132–141 (2011).

65. A. M. Sanchez, R. Candau, H. Bernardi, Recent Data on Cellular Component Turnover: Focus on Adaptations to Physical Exercise. Cells 8 (2019).

66. A. M. Sanchez et al., The role of AMP-activated protein kinase in the coordination of skeletal muscle turnover and energy homeostasis. Am J Physiol Cell Physiol 303, C475–485 (2012).

67. D. F. Egan et al., Phosphorylation of ULK1 (hATG1) by AMP-activated protein kinase connects energy sensing to mitophagy. Science 331, 456–461 (2011).

68. C. G. Pack et al., Quantitative live-cell imaging reveals spatio-temporal dynamics and cytoplasmic assembly of the 26S proteasome. Nat Commun 5, 3396 (2014).

69. S. Asano et al., Proteasomes. A molecular census of 26S proteasomes in intact neurons. Science 347, 439–442 (2015).

70. T. Miyamoto et al., Compartmentalized AMPK signaling illuminated by genetically encoded molecular sensors and actuators. Cell Rep 11, 657–670 (2015).

71. D. L. Schmitt et al., Spatial regulation of AMPK signaling revealed by a sensitive kinase activity reporter. Nat Commun 13, 3856 (2022).

72. J. C. Drake et al., Mitochondria-localized AMPK responds to local energetics and contributes to exercise and energetic stress-induced mitophagy. Proc Natl Acad Sci U S A 118 (2021).

73. K. Yamano, A. I. Fogel, C. Wang, A. M. van der Bliek, R. J. Youle, Mitochondrial Rab GAPs govern autophagosome biogenesis during mitophagy. Elife 3, e01612 (2014).

74. A. Rak et al., Crystal structure of the GAP domain of Gyp1p: first insights into interaction with Ypt/Rab proteins. EMBO J 19, 5105–5113 (2000).

75. S. Han, Y. Y. Jeong, P. Sheshadri, X. Su, Q. Cai, Mitophagy regulates integrity of mitochondria at synapses and is critical for synaptic maintenance. EMBO Rep 10.15252/embr.201949801, e201949801 (2020).

76. A. H. Nagahara et al., Early BDNF treatment ameliorates cell loss in the entorhinal cortex of APP transgenic mice. J Neurosci 33, 15596–15602 (2013).

77. V. A. Polito et al., Selective clearance of aberrant tau proteins and rescue of neurotoxicity by transcription factor EB. EMBO Mol Med 6, 1142–1160 (2014).

78. X. Ye et al., Regulation of Synaptic Amyloid-beta Generation through BACE1 Retrograde Transport in a Mouse Model of Alzheimer’s Disease. J Neurosci 37, 2639–2655 (2017).

79. A. H. Nagahara et al., Neuroprotective effects of brain-derived neurotrophic factor in rodent and primate models of Alzheimer’s disease. Nat Med 15, 331–337 (2009).

80. Q. Xiao et al., Neuronal-Targeted TFEB Accelerates Lysosomal Degradation of APP, Reducing Abeta Generation and Amyloid Plaque Pathogenesis. J Neurosci 35, 12137–12151 (2015).

81. Y. Xie et al., Endolysosomal Deficits Augment Mitochondria Pathology in Spinal Motor Neurons of Asymptomatic fALS Mice. Neuron 87, 355–370 (2015).

82. S. G. Greenberg, P. Davies, A preparation of Alzheimer paired helical filaments that displays distinct tau proteins by polyacrylamide gel electrophoresis. Proc Natl Acad Sci U S A 87, 5827–5831 (1990).

83. J. Lewis et al., Neurofibrillary tangles, amyotrophy and progressive motor disturbance in mice expressing mutant (P301L) tau protein. Nat Genet 25, 402–405 (2000).

84. N. Sahara et al., Assembly of tau in transgenic animals expressing P301L tau: alteration of phosphorylation and solubility. J Neurochem 83, 1498–1508 (2002).

85. N. Sahara et al., Characteristics of TBS-extractable hyperphosphorylated tau species: aggregation intermediates in rTg4510 mouse brain. J Alzheimers Dis 33, 249–263 (2013).

86. Y. Ren, N. Sahara, Characteristics of tau oligomers. Front Neurol 4, 102 (2013).

87. J. N. Crawley, Designing mouse behavioral tasks relevant to autistic-like behaviors. Ment Retard Dev Disabil Res Rev 10, 248–258 (2004).

88. S. S. Moy et al., Sociability and preference for social novelty in five inbred strains: an approach to assess autistic-like behavior in mice. Genes Brain Behav 3, 287–302 (2004).

89. J. J. Nadler et al., Automated apparatus for quantitation of social approach behaviors in mice. Genes Brain Behav 3, 303–314 (2004).

90. H. Takeuchi et al., P301S mutant human tau transgenic mice manifest early symptoms of human tauopathies with dementia and altered sensorimotor gating. PLoS One 6, e21050 (2011).

91. J. Nunez, Morris Water Maze Experiment. J Vis Exp 10.3791/897 (2008).

92. R. Glass, S. Norton, N. Fox, A. W. Kusnecov, Maternal immune activation with staphylococcal enterotoxin A produces unique behavioral changes in C57BL/6 mouse offspring. Brain Behav Immun 75, 12–25 (2019).

93. R. G. Phillips, J. E. LeDoux, Differential contribution of amygdala and hippocampus to cued and contextual fear conditioning. Behav Neurosci 106, 274–285 (1992).

94. A. Litvinchuk et al., Complement C3aR Inactivation Attenuates Tau Pathology and Reverses an Immune Network Deregulated in Tauopathy Models and Alzheimer’s Disease. Neuron 100, 1337–1353 e1335 (2018).

95. W. Peng, Y. C. Wong, D. Krainc, Mitochondria-lysosome contacts regulate mitochondrial Ca(2+) dynamics via lysosomal TRPML1. Proc Natl Acad Sci U S A 117, 19266–19275 (2020).

96. Q. Cai, D. Ganesan, Regulation of neuronal autophagy and the implications in neurodegenerative diseases. Neurobiol Dis 162, 105582 (2022).

97. R. A. Nixon, The role of autophagy in neurodegenerative disease. Nat Med 19, 983–997 (2013).

98. A. K. H. Stavoe, E. L. F. Holzbaur, Autophagy in Neurons. Annu Rev Cell Dev Biol 35, 477–500 (2019).

99. D. K. Sidibe, M. C. Vogel, S. Maday, Organization of the autophagy pathway in neurons. Curr Opin Neurobiol 75, 102554 (2022).

100. N. C. Chan et al., Broad activation of the ubiquitin-proteasome system by Parkin is critical for mitophagy. Hum Mol Genet 20, 1726–1737 (2011).

101. S. R. Yoshii, C. Kishi, N. Ishihara, N. Mizushima, Parkin mediates proteasome-dependent protein degradation and rupture of the outer mitochondrial membrane. J Biol Chem 286, 19630–19640 (2011).

102. D. Popovic et al., Rab GTPase-activating proteins in autophagy: regulation of endocytic and autophagy pathways by direct binding to human ATG8 modifiers. Mol Cell Biol 32, 1733–1744 (2012).

103. A. Bhattacharya et al., A lysosome membrane regeneration pathway depends on TBC1D15 and autophagic lysosomal reformation proteins. Nat Cell Biol 25, 685–698 (2023).

104. K. Yamano et al., Endosomal Rab cycles regulate Parkin-mediated mitophagy. Elife 7 (2018).

105. B. R. Yan et al., C5orf51 is a component of the MON1-CCZ1 complex and controls RAB7A localization and stability during mitophagy. Autophagy 18, 829–840 (2022).

106. X. Jin et al., RAB7 activity is required for the regulation of mitophagy in oocyte meiosis and oocyte quality control during ovarian aging. Autophagy 18, 643–660 (2022).

107. A. Jimenez-Orgaz et al., Control of RAB7 activity and localization through the retromer-TBC1D5 complex enables RAB7-dependent mitophagy. EMBO J 37, 235–254 (2018).

108. W. Yu et al., TBC1D15/RAB7-regulated mitochondria-lysosome interaction confers cardioprotection against acute myocardial infarction-induced cardiac injury. Theranostics 10, 11244–11263 (2020).

109. S. Sun et al., TBC1D15-Drp1 interaction-mediated mitochondrial homeostasis confers cardioprotection against myocardial ischemia/reperfusion injury. Metabolism 134, 155239 (2022).

110. S. Camandola, M. P. Mattson, Brain metabolism in health, aging, and neurodegeneration. EMBO J 36, 1474–1492 (2017).

111. K. Banerjee, S. Munshi, D. E. Frank, G. E. Gibson, Abnormal Glucose Metabolism in Alzheimer’s Disease: Relation to Autophagy/Mitophagy and Therapeutic Approaches. Neurochem Res 40, 2557–2569 (2015).

112. B. J. Neth, S. Craft, Insulin Resistance and Alzheimer’s Disease: Bioenergetic Linkages. Front Aging Neurosci 9, 345 (2017).

113. E. Tonnies, E. Trushina, Oxidative Stress, Synaptic Dysfunction, and Alzheimer’s Disease. J Alzheimers Dis 57, 1105–1121 (2017).

114. M. F. Beal, Mitochondria take center stage in aging and neurodegeneration. Ann Neurol 58, 495–505 (2005).

115. P. Bubber, V. Haroutunian, G. Fisch, J. P. Blass, G. E. Gibson, Mitochondrial abnormalities in Alzheimer brain: mechanistic implications. Ann Neurol 57, 695–703 (2005).

116. R. A. Nixon, D. C. Rubinsztein, Mechanisms of autophagy-lysosome dysfunction in neurodegenerative diseases. Nat Rev Mol Cell Biol 25, 926–946 (2024).

117. F. Lim et al., FTDP-17 mutations in tau transgenic mice provoke lysosomal abnormalities and Tau filaments in forebrain. Mol Cell Neurosci 18, 702–714 (2001).

118. N. Cummins, A. Tweedie, S. Zuryn, J. Bertran-Gonzalez, J. Gotz, Disease-associated tau impairs mitophagy by inhibiting Parkin translocation to mitochondria. EMBO J 38 (2019).

119. E. F. Fang et al., Mitophagy inhibits amyloid-beta and tau pathology and reverses cognitive deficits in models of Alzheimer’s disease. Nat Neurosci 22, 401–412 (2019).

120. S. Maday, K. E. Wallace, E. L. Holzbaur, Autophagosomes initiate distally and mature during transport toward the cell soma in primary neurons. J Cell Biol 196, 407–417 (2012).

121. S. E. Hill et al., Maturation and Clearance of Autophagosomes in Neurons Depends on a Specific Cysteine Protease Isoform, ATG-4.2. Dev Cell 49, 251–266 e258 (2019).

122. N. D. Okerlund et al., Bassoon Controls Presynaptic Autophagy through Atg5. Neuron 93, 897–913 e897 (2017).

123. S. F. Soukup et al., A LRRK2-Dependent EndophilinA Phosphoswitch Is Critical for Macroautophagy at Presynaptic Terminals. Neuron 92, 829–844 (2016).

124. J. Yamaguchi et al., Atg9a deficiency causes axon-specific lesions including neuronal circuit dysgenesis. Autophagy 14, 764–777 (2018).

125. P. P. Y. Lie et al., Axonal transport of late endosomes and amphisomes is selectively modulated by local Ca(2+) efflux and disrupted by PSEN1 loss of function. Sci Adv 8, eabj5716 (2022).

126. C. Giorgi, S. Marchi, P. Pinton, The machineries, regulation and cellular functions of mitochondrial calcium. Nat Rev Mol Cell Biol 19, 713–730 (2018).

127. M. Kumar et al., Triglycerides are an important fuel reserve for synapse function in the brain. Nat Metab 7, 1392–1403 (2025).

128. S. H. Saber et al., DDHD2 provides a flux of saturated fatty acids for neuronal energy and function. Nat Metab 7, 2117–2141 (2025).

129. V. Nikoletopoulou, N. Tavernarakis, Regulation and Roles of Autophagy at Synapses. Trends Cell Biol 28, 646–661 (2018).

130. A. Karpova et al., Neuronal autophagy in the control of synapse function. Neuron 113, 974–990 (2025).

131. C. Marinangeli et al., AMP-Activated Protein Kinase Is Essential for the Maintenance of Energy Levels during Synaptic Activation. iScience 9, 1–13 (2018).

132. Y. Yoshiyama et al., Synapse loss and microglial activation precede tangles in a P301S tauopathy mouse model. Neuron 53, 337–351 (2007).

133. S. Han, M. Zhang, Y. Y. Jeong, D. J. Margolis, Q. Cai, The role of mitophagy in the regulation of mitochondrial energetic status in neurons. Autophagy 10.1080/15548627.2021.1907167, 1-20 (2021).

134. Q. Cai et al., Snapin-regulated late endosomal transport is critical for efficient autophagy-lysosomal function in neurons. Neuron 68, 73–86 (2010).

135. Q. Cai, H. M. Zakaria, A. Simone, Z. H. Sheng, Spatial parkin translocation and degradation of damaged mitochondria via mitophagy in live cortical neurons. Curr Biol 22, 545–552 (2012).

136. X. Ye, Q. Cai, Snapin-mediated BACE1 retrograde transport is essential for its degradation in lysosomes and regulation of APP processing in neurons. Cell Rep 6, 24–31 (2014).

137. X. Ye, X. Sun, V. Starovoytov, Q. Cai, Parkin-mediated mitophagy in mutant hAPP neurons and Alzheimer’s disease patient brains. Hum Mol Genet 24, 2938–2951 (2015).

138. G. A. Banker, W. M. Cowan, Further observations on hippocampal neurons in dispersed cell culture. J Comp Neurol 187, 469–493 (1979).

139. J. S. Kang et al., Docking of axonal mitochondria by syntaphilin controls their mobility and affects short-term facilitation. Cell 132, 137–148 (2008).

140. D. N. Mastronarde, S. R. Held, Automated tilt series alignment and tomographic reconstruction in IMOD. J Struct Biol 197, 102–113 (2017).

141. A. Martinez-Sanchez, I. Garcia, S. Asano, V. Lucic, J. J. Fernandez, Robust membrane detection based on tensor voting for electron tomography. J Struct Biol 186, 49–61 (2014).

142. E. F. Pettersen et al., UCSF Chimera--a visualization system for exploratory research and analysis. J Comput Chem 25, 1605–1612 (2004).

143. R. A. Nixon et al., Extensive involvement of autophagy in Alzheimer disease: an immuno-electron microscopy study. J Neuropathol Exp Neurol 64, 113–122 (2005).

144. D. J. Klionsky et al., Guidelines for the use and interpretation of assays for monitoring autophagy (3rd edition). Autophagy 12, 1–222 (2016).

145. L. Scorrano et al., Coming together to define membrane contact sites. Nat Commun 10, 1287 (2019).

146. Z. Yu et al., Beyond t test and ANOVA: applications of mixed-effects models for more rigorous statistical analysis in neuroscience research. Neuron 110, 21–35 (2022).

