## Supplemental information for "Dysregulation of a novel autophagosome-mitochondria contact contributes to autophagy dysfunction and neurodegeneration in tauopathy"

### Figures

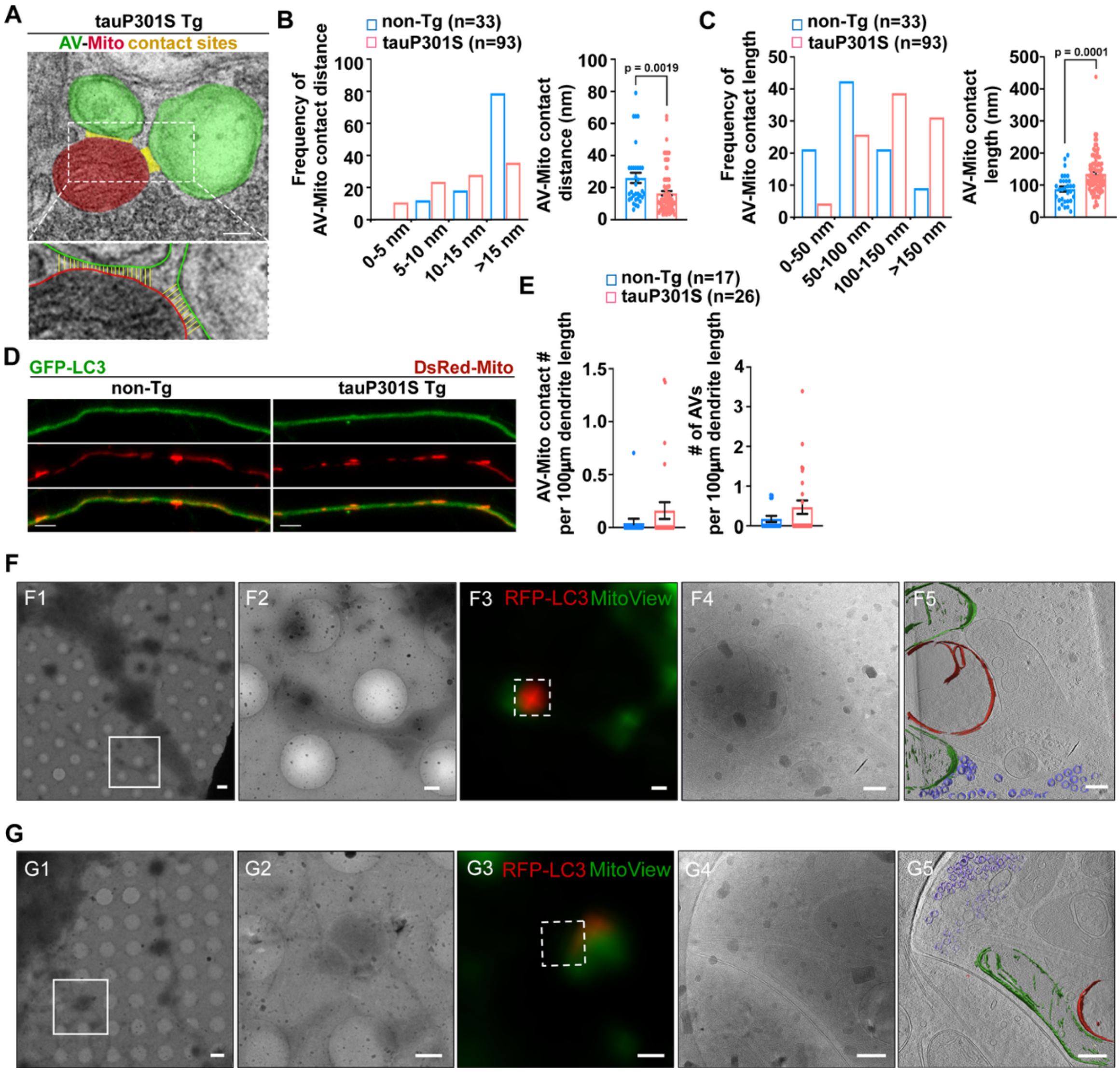


**Fig. S1.** Autophagosome/autophagic vacuole (AV)-mitochondria (Mito) contacts in tauopathy neurons. (*A*-*C*) The representative pseudocolored transmission electron microscopy (TEM) images from tauP301S Tg (PS19) mouse brains highlighting the AVs (green), the mitochondrion (red), and AV-Mito contact sites (yellow lines) (*A*). AV-Mito contact sites are defined as sites of contact within a reciprocal distance of 80 nm. The bottom panel shows a zoomed-in view of a selected area (marked with a white-dashed box) in the corresponding panel. Yellow lines indicate the AV-Mito contacts. The intermembrane distance and length of individual AV-Mito contact sites, as well as their frequencies, were quantified and compared to those in non-Tg mouse brains (*B* and *C*). Data were collected from three pairs of non-Tg and tauP301S Tg mice: the total numbers of AV-Mito contacts (*n*) are indicated in parentheses (*B* and *C*). (*D* and *E*) Representative images

(*D*) and quantitative analysis (*E*) of AV and AV-Mito contact densities in the dendritic processes of non-Tg and tauP301S Tg neurons. Non-Tg and tauP301S Tg neurons were imaged after 24-hour incubation with 100 mM trehalose. The data were expressed as the number of AV-Mito contacts or AVs per 100 μm axonal length in non-Tg or tauP301S Tg neurons, respectively. Data were collected from two dissections: the total numbers of neurons (*n*) are indicated in parentheses (*E*). (*F* and *G*) Correlative light and electron microscope (CLEM) and cryo-electron tomography (cryo-ET) of tauP301S Tg neurons showing AV-Mito contacts in axonal processes. Low-magnification cryo-EM images (740 ×) of the neuronal process region containing AV-Mito contacts (*F1* and *G1*). Cryo-EM image of the boxed area in *F1* and *G1*—acquired at 3,600 × magnification to further localize candidate AV-Mito contact sites (*F2* and *G2*). Cryo-fluorescence image of the same regions—highlighting putative RFP-LC3-marked AVs and MitoView Green-labeled mitochondria. These images were used as references and correlated with the 3,600 × cryo-EM images to precisely target AV-Mito contact sites (*F3* and *G3*). The 0° tilt image of the tilt series taken in the regions outlined by the dashed box in *F3* and *G3*, acquired at 26,000 × magnification (*F4* and *G4*). Annotated views of the tomogram depicting the mitochondrion (green), the AV (red), and synaptic vesicles (purple) on the background of the central slice of the tomogram (*F5* and *G5*). TauP301S Tg neurons were treated with 80 mM trehalose at DIV13 prior to imaging. Data were expressed as the mean ± SEM and analyzed using linear mixed-effects models (*B* and *C*) or by two-sided unpaired Student's *t*-test (*E*). Scale bars: 100 nm (*A*), 10 μm (*D*), 2 μm (*F1* and *G1*), 1 μm (*F2, F3, G2,* and *G3*), and 200 nm (*F4, F5, G4,* and *G5*).

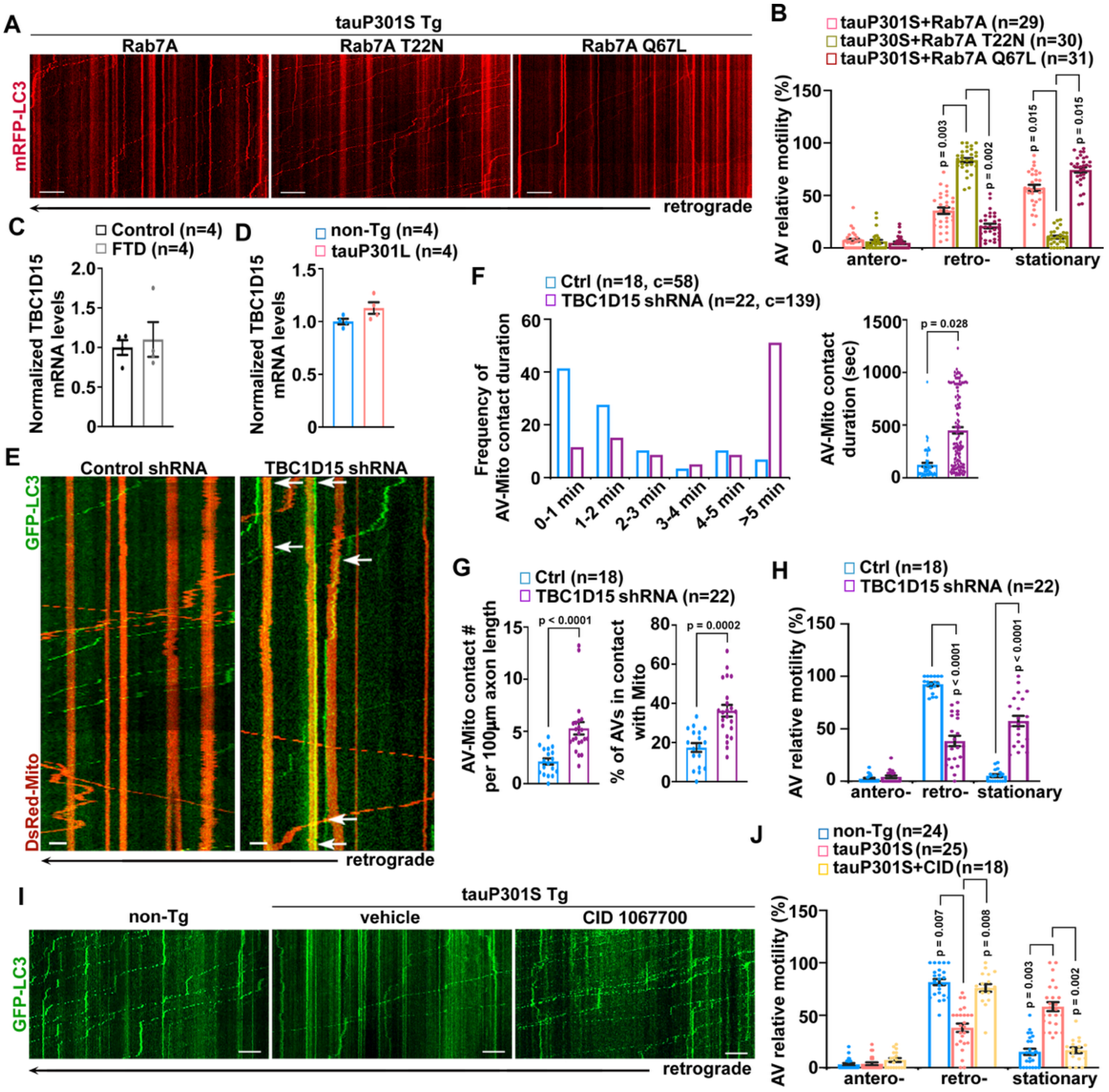


**Fig. S2.** Rab7 GAP TBC1D15 deficiency leads to defects in AV-Mito contact untethering and AV retrograde transport in axons. (*A* and *B*) Representative kymographs (A) and quantitative analysis (*B*) of AV motility in tauP301S Tg axons expressing Rab7A, Rab7A T22N, or Rab7A Q67L. Relative motility of AVs was quantified in these tauP301S Tg axons. Vertical lines represent stationary organelles; slanted lines to the right (negative slope) represent anterograde movement; those to the left (positive slope) indicate retrograde movement. An AV was considered stationary if it remained immotile (displacement ≤ 5 μm). Data were collected from two dissections: the total numbers of neurons (*n*) are indicated in parentheses (*B*). (*C*) TBC1D15 mRNA levels are unaltered in FTD patient brains. TBC1D15 mRNA levels were normalized to GAPDH and to those of normal control subjects. (*D*) TauP301L Tg mouse brains show no detectable change in TBC1D15 mRNA levels. TBC1D15 mRNA levels were normalized to GAPDH and to those of non-

Tg littermate controls. (*E-H*) Representative kymographs (*E*) and quantitative analysis (*F-H*) of AV-Mito contact dynamics in the axons of non-Tg neurons expressing TBC1D15 shRNA. The data were quantified and expressed as the frequency of AV-Mito contact duration, the average duration of AV-Mito contacts, the number of AV-Mito contacts per 100 μm axonal length, the percentage of AVs in AV-Mito contacts, and the relative motility of AVs in the axons of non-Tg neurons with and without TBC1D15 RNAi in the presence of trehalose. Arrows: AV-Mito contacts. Data were collected from two dissections: the total numbers of neurons (*n*) are indicated in parentheses (*F-H*). (*I* and *J*) Representative kymographs (I) and quantitative analysis (J) of AV motility in the axons of non-Tg neurons and tauP301S Tg neurons with and without CID 1067700 (80 nM) treatment. The data were collected from two dissections and expressed as the relative motility of AVs in axons from the total numbers of neurons (*n*) as indicated in parentheses (*J*). Data were expressed as the mean ± SEM and analyzed using linear mixed-effects models (*B*, *F-H* and *J*) or by two-sided unpaired Student's *t*-test (*C*, *D*). Scale bars: 10 μm.

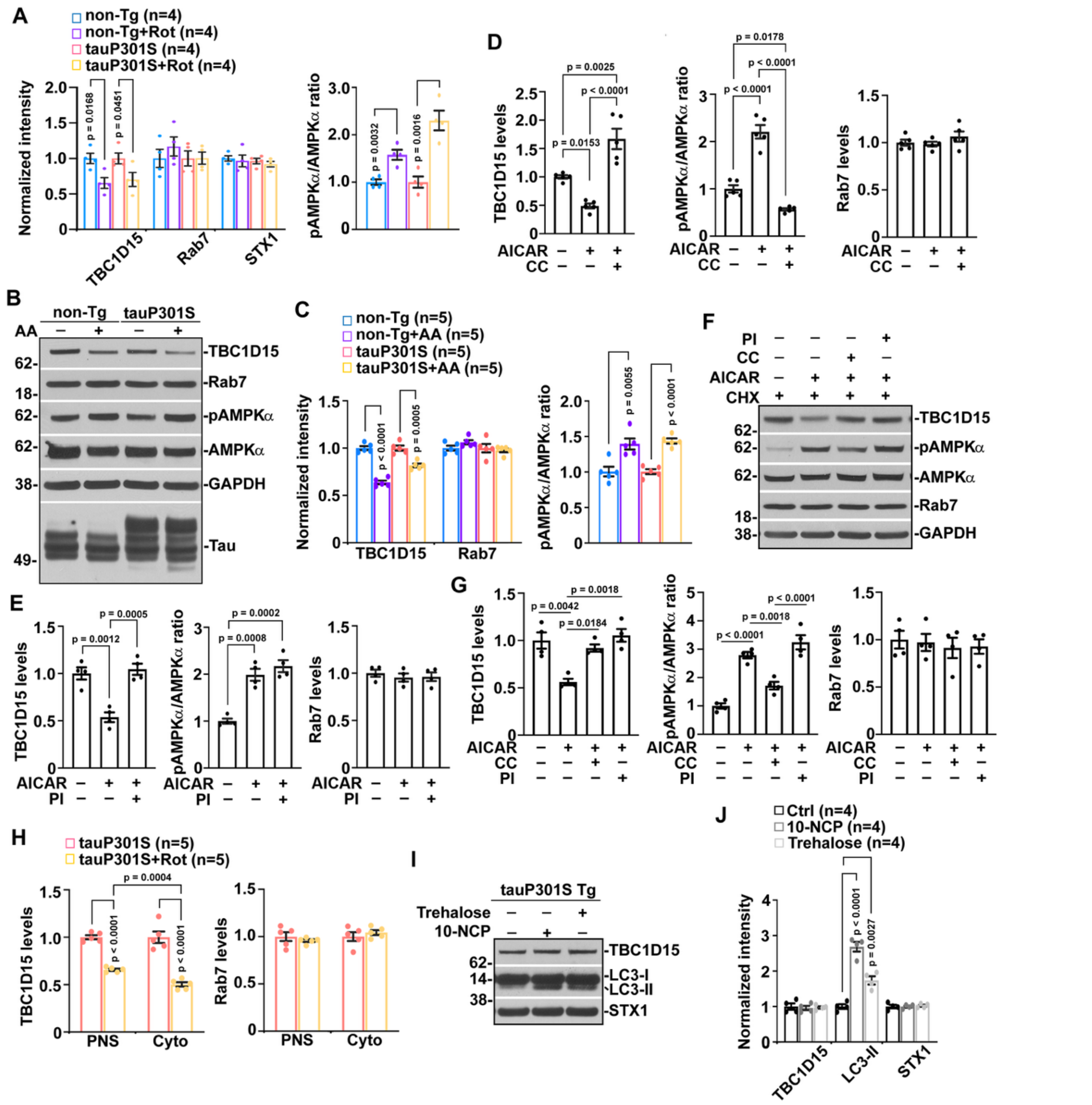


**Fig. S3.** Increased AMPK activity accelerates TBC1D15 turnover in bioenergetically stressed neurons. (*A*) Quantitative analysis of non-Tg and tauP301S Tg neurons with and without 24-hour incubation of rotenone (20 μM). Protein intensities were normalized to those in vehicle-treated control non-Tg or tauP301S Tg neurons. Rot: rotenone. (*B* and *C*) Representative blots (*B*) and quantitative analysis (*C*) showing decreases in TBC1D15 levels in non-Tg and in tauP301S Tg neurons treated with antimycin A (5 nM) for 24 hours. Protein intensities were normalized to those in vehicle-treated control non-Tg or tauP301S Tg neurons. Data were quantified from four dissections. AA: antimycin A. (*D*) Quantitative analysis of tauP301S Tg neurons incubated with AICAR (0.5 mM) or/and CC (5 μM) in the presence of cycloheximide (10 μg/ml) for 24 hours. Protein intensities were normalized to those of tauP301S controls. AICAR: 5-aminoimidazole-4-carboxamide riboside; CC: Compound C. (*E*) Quantitative analysis of tauP301S Tg neurons treated with and without AICAR (0.5 mM) and/or epoxomicin (100 nM), a proteasome inhibitor, in the presence of cycloheximide (10 μg/ml). Protein intensities were normalized to those of tauP301S controls. PI: proteasome inhibitor. (*F* and *G*) Representative blots (*F*) and quantitative analysis (*G*) of TBC1D15 levels in neurons with and without 24-hour 1 mM AICAR treatment in the presence and absence of 1 μM CC or epoxomicin (100 nM). CHX: cycloheximide (10 μg/ml). Protein intensities were quantified and normalized to those of control neurons. (*H*) Quantitative analysis of total and cytosolic TBC1D15 levels in tauP301S Tg neurons with and without rotenone (20 μM). Total and cytosolic TBC1D15 levels were normalized to those in tauP301S controls without rotenone treatment. PNS: post-nuclear supernatant; Cyto: cytosolic fraction. (*I* and *J*) Representative blots (*I*) and quantitative analysis (*J*) of TBC1D15 expression levels in tauP301S Tg neurons with and without 24-hour trehalose (100 mM) or 10-NCP (10 μM) incubation. Protein intensities were normalized to those in vehicle-treated control tauP301S Tg neurons. Data were collected from four (*A, E, G,* and *J*) or five (*C*, *D,* and *H*) dissections. Data were expressed as the mean ± SEM and analyzed by two-sided unpaired Student's *t*-test (*A, C,* and *H*) or one-way ANOVA with Tukey's correction (*D, E, G,* and *J*).

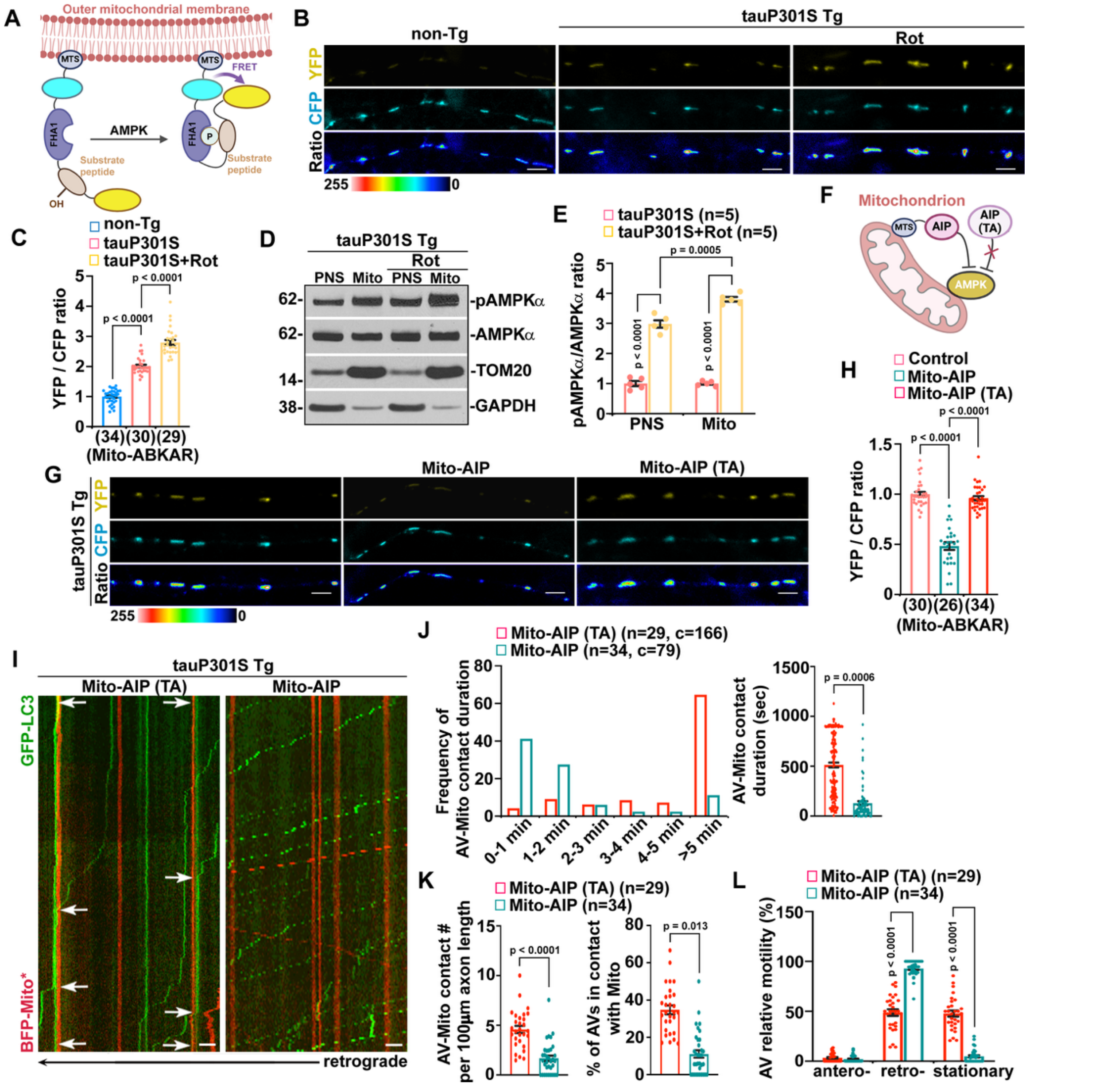


**Fig. S4.** AV-Mito contact release defects are attributed to the hyperactivity of mitochondria-localized AMPK in tauopathy neurons. (*A*) Schematic diagram of Mito-ABKAR, a highly sensitive FRET-based mitochondrial AMPK reporter. Mito-ABKAR consists of MTS, CFP variant, Cerulean (cyan), FHA1 phospho-amino acid-binding domain, AMPK substrate motif/peptide, and YFP variant cpVE172 (yellow). The FRET signal on mitochondria exhibits an AMPK-dependent pattern through the AMPK phosphorylation of the substrate peptide. FRET: Förster resonance energy transfer; MTS: mitochondria-targeting sequence**.** (*B* and *C*) Representative images (*B*) and quantitative analysis (*C*) of mitochondrial AMPK activity in non-Tg and tauP301S Tg neurons. The YFP/CFP emission ratio of Mito-ABKAR on axonal mitochondria was measured and normalized to that in non-Tg neurons without rotenone treatment. Data were quantified from the total numbers of neurons (*n*) as indicated in parentheses (*C*). (*D* and *E*) Representative blots (*D*) and quantitative analysis (*E*) of AMPK activity in tauP301S Tg neurons with and without 24-hour rotenone treatment. The pAMPK$\alpha$/AMPK$\alpha$ ratio in PNS and mitochondrial fractions was normalized to that in control tauP301S Tg neurons. PNS: post-nuclear supernatant; Mito: mitochondrial fraction. (*F*) Schematic diagram of Mito-AIP and Mito-AIP (TA). Mito-AIP is composed of MTS and AIP or AIP (TA) mutant showing no effect on inhibition of mitochondria-localized AMPK activity. AIP: AMPK inhibitor peptide. (*G* and *H*) Representative images (*G*) and quantitative analysis (*H*) of mitochondrial AMPK activity in tauP301S Tg axons expressing mitochondrial AMPK inhibitor peptide (Mito-AIP), but not Mito-AIP (TA), which cannot be phosphorylated by AMPK. The YFP/CFP emission ratio of Mito-ABKAR on axonal mitochondria was measured and normalized to that in control tauP301S Tg neurons. Data were quantified from the total numbers of neurons (*n*) as indicated in parentheses (*H*). (*I*-*L*) Representative kymographs (*I*) and quantitative analysis (*J*-*L*) of AV-Mito contacts and AVs in tauP301S Tg axons expressing Mito-AIP (TA) or Mito-AIP. The data were quantified and expressed as the duration of AV-Mito contacts, the frequency of AV-Mito contact duration, the number of AV-Mito contacts per 100 $\mu$m axonal length, the percentage of AVs in AV-Mito contacts, and the relative motility of AVs in Mito-AIP-expressing tauP301S axons, compared to those of tauP301S controls expressing Mito-AIP (TA). Arrows: AV-Mito contacts. Data were quantified from the total numbers of neurons (*n*) and AV-Mito contacts (*c*) as indicated in parentheses (*J*-*L*). Data were collected from three (*C*, *H*, and *J*-*L*) or five (*E*) dissections. Data were expressed as the mean $\pm$ SEM and analyzed using linear mixed-effects models (*C, H,* and *J*-*L*) or by two-sided unpaired Student's *t*-test (*E*). Scale bars: 10 $\mu$m.

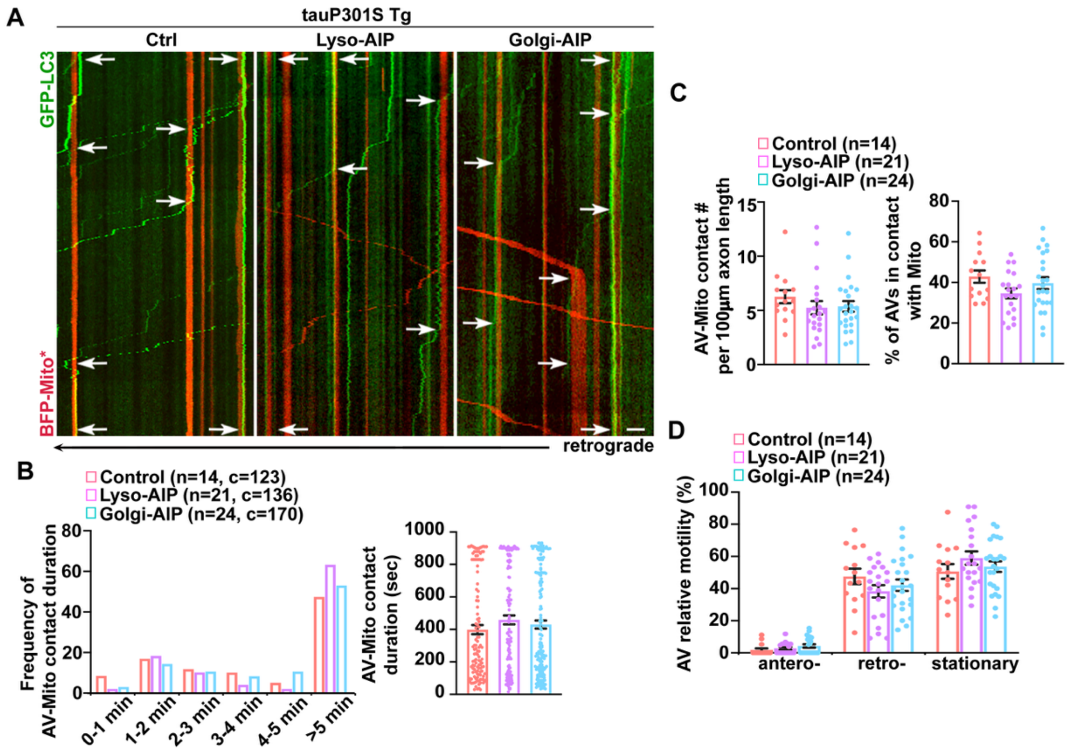


**Fig. S5.** Inhibition of lysosome or Golgi-localized AMPK activity fails to reverse defective AV-Mito untethering in tauopathy axons. (*A*-*D*) Representative kymographs (*A*) and quantitative analysis (*B*-*D*) of AV-Mito contact dynamics in tauP301S Tg axons expressing lyso-AIP or Golgi-AIP after 24-hour trehalose treatment. The data were quantified and expressed as the duration of AV-Mito contacts, the frequency of AV-Mito contact duration, the number of AV-Mito contacts per 100 μm axonal length, the percentage of AVs in AV-Mito contacts, and the relative motility of AVs in the axons, compared to those of control tauP301S Tg neurons. Data were collected from three dissections: the total numbers of neurons (*n*) and AV-Mito contacts (*c*) are indicated in parentheses (*B*-*D*). Arrows: AV-Mito contacts. Data were expressed as the mean ± SEM and analyzed using linear mixed-effects models. Scale bars: 10 μm.

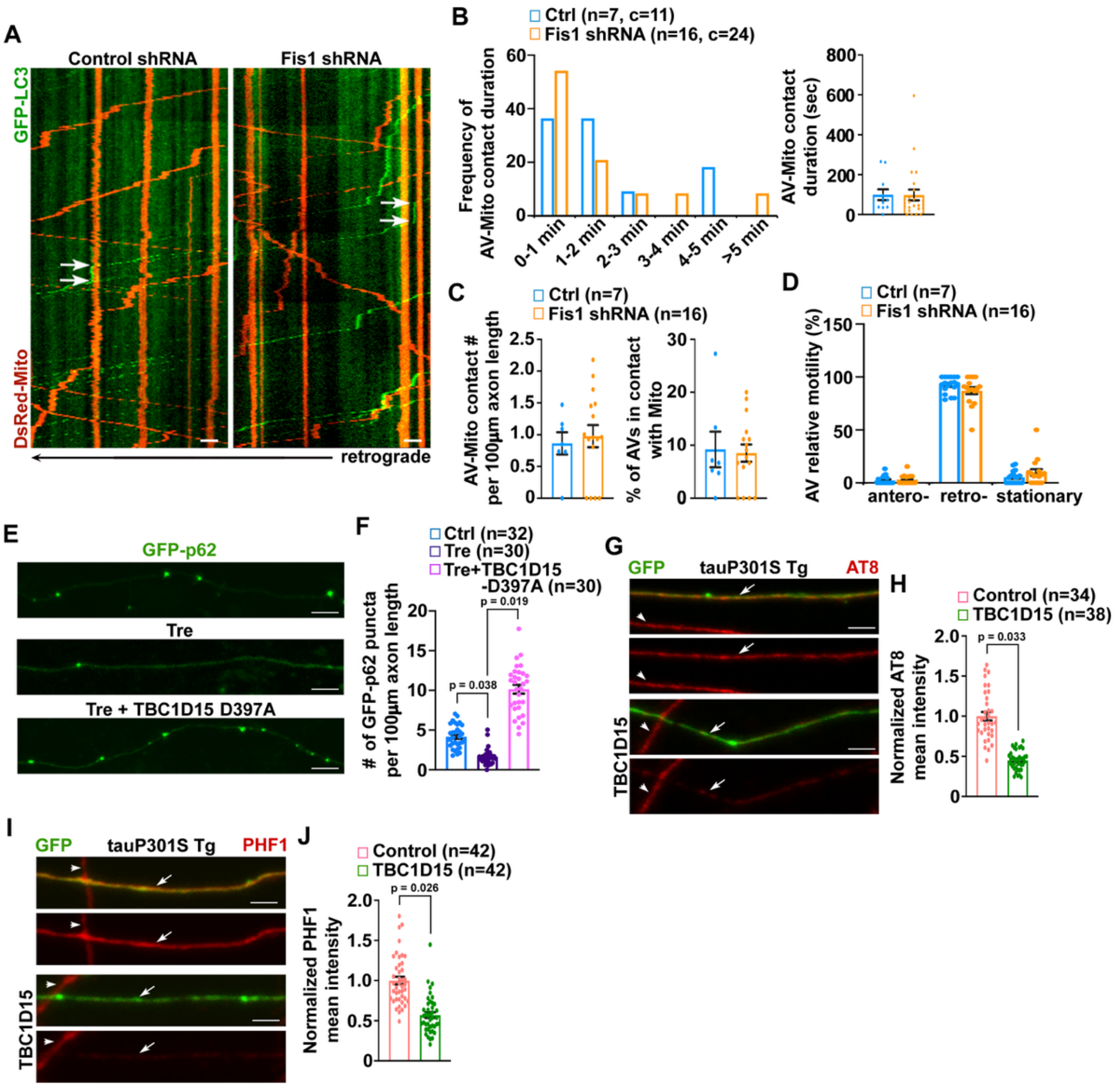

**Fig. S6.** AVs recruit non-mitochondrial TBC1D15 that releases AV-Mito contacts in axons. (*A-D*) Representative kymographs (*A*) and quantitative analysis (*B-D*) of AV-Mito contact dynamics in non-Tg axons expressing Fis1 shRNA. The data were quantified and expressed as the duration of AV-Mito contacts, the frequency of AV-Mito contact duration, the number of AV-Mito contacts per 100 μm axonal length, the percentage of AVs in AV-Mito contacts, and the relative motility of AVs in the axons of non-Tg neurons with and without loss of Fis1 in the presence of trehalose. Data were collected from two dissections: the total numbers of neurons (*n*) and AV-Mito contacts (c) are indicated in parentheses (*B-D*). Arrows: AV-Mito contacts. (*E* and *F*) Representative images (*E*) and quantitative analysis (*F*) of autophagic substrate p62 in axons with and without overexpression (OE) of TBC1D15 D397A mutant or trehalose treatment. The data were quantified and expressed as the number of GFP-p62 puncta per 100 μm axonal length in neurons. Data were collected from three dissections: the total numbers of neurons (*n*) are indicated in parentheses (*F*). (*G*-*J*) Representative images (*G* and *I*) and quantitative analysis (*H* and *J*) of phospho-tau levels in the axons of tauP301S Tg neurons with and without OE of TBC1D15. Phospho-tau was visualized by AT8 and PHF1 antibodies. Arrows: axons expressing GFP or GFP with TBC1D15; arrowheads: untransfected axons in the same imaging field. The mean intensities of AT8 or PHF1-marked phospho-tau were normalized to those of control tauP301S Tg axons. Data were collected from three dissections: the total numbers of neurons (*n*) are indicated in parentheses (*H* and *J*). Data were expressed as the mean ± SEM and analyzed using linear mixed-effects models. Scale bars: 10 μm.

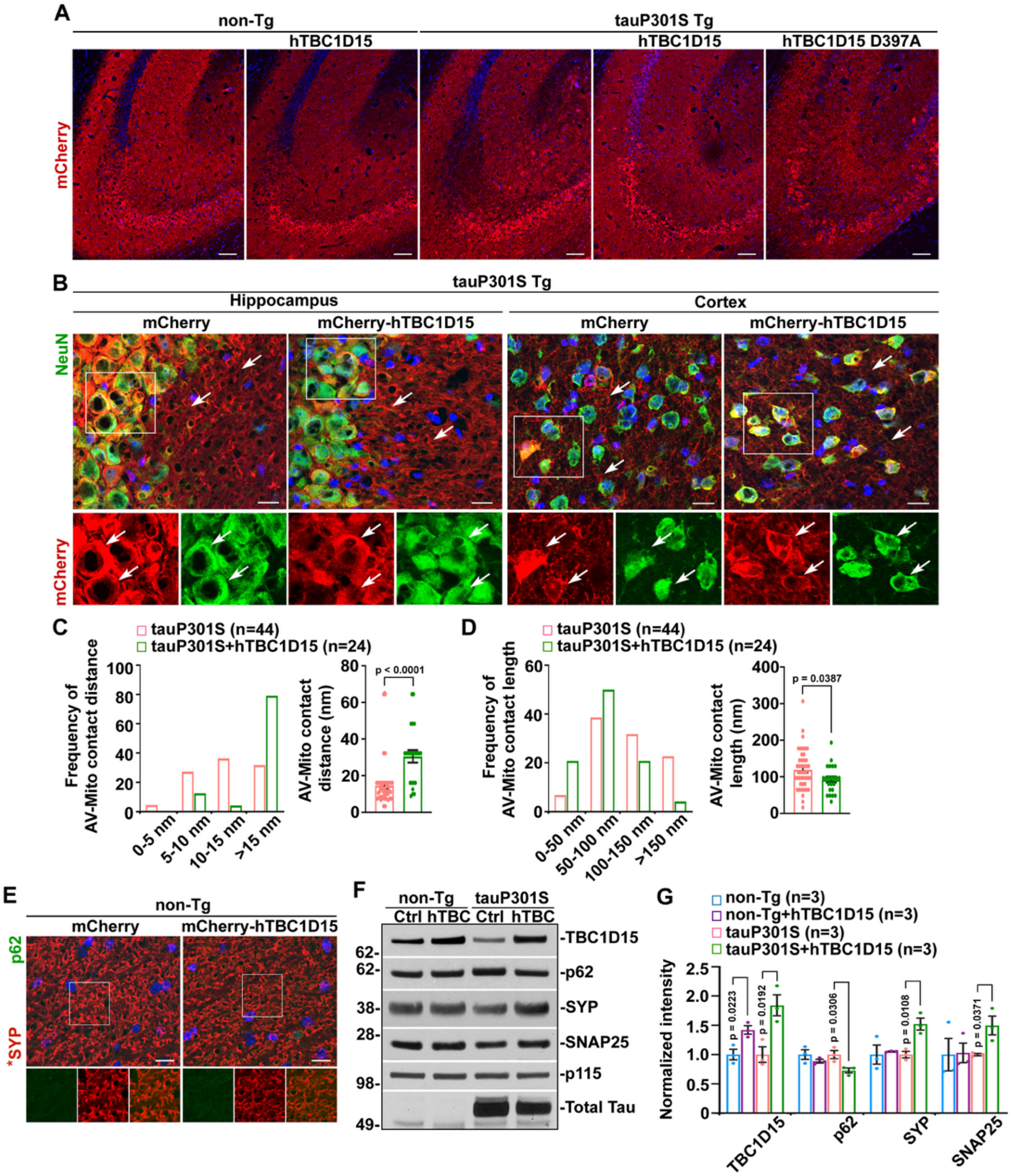


**Fig. S7.** The rescue effects of AAV injection-mediated TBC1D15 OE in tauopathy mouse brains. (*A*) Representative images showing gene delivery into the hippocampus of non-Tg or tauP301S Tg mice injected with AAV-mCherry, AAV-mCherry-hTBC1D15, or AAV-mCherry-hTBC1D15 D397A. mCherry fluorescence is present in transduced neurons in the hippocampal regions of non-Tg and tauP301S Tg mouse brains. (*B*) Representative images showing mCherry fluorescence in the soma and processes (arrows) of hippocampal and cortical neurons in the brains of AAV-mCherry or AAV-mCherry-hTBC1D15-injected tauP301S Tg mice at 8 months of age. (*C* and *D*) Quantitative analysis of TEM images showing increased distance but reduced length of AV-Mito contact structures in the hippocampal regions of 8-month-old tauP301S Tg mouse brains injected with AAV-mCherry-hTBC1D15. The distance and length of individual AV-Mito contact sites, as well as their frequencies, were quantified and compared to those in the control AAV-mCherry-infected tauP301S Tg mouse brains. Data were quantified from a total number of AV-Mito contacts (*n*) as indicated in parentheses (*C* and *D*) from two mice per group. (*E*) Representative images showing p62 fluorescence in the hippocampus of 10-month-old non-Tg mouse brains with and without TBC1D15 OE. SYP: synaptophysin. (*F* and *G*) Representative blots (*F*) and quantitative analysis (*G*) showing that TBC1D15 OE led to a decrease in p62 but increases in the levels of synaptic protein markers: SYP and SNAP25 in the hippocampi of 10-month-old tauP301S Tg mouse brains. Protein intensities were normalized to those in the control non-Tg mice or tauP301S Tg mice without TBC1D15 OE. Data were quantified from three pairs of mice. Data were expressed as the mean ± SEM and analyzed using linear mixed-effects models (*C* and *D*) or by two-sided unpaired Student's *t*-test (*G*). Scale bars: 250 μm (*A*) and 10 μm (*B* and *E).*

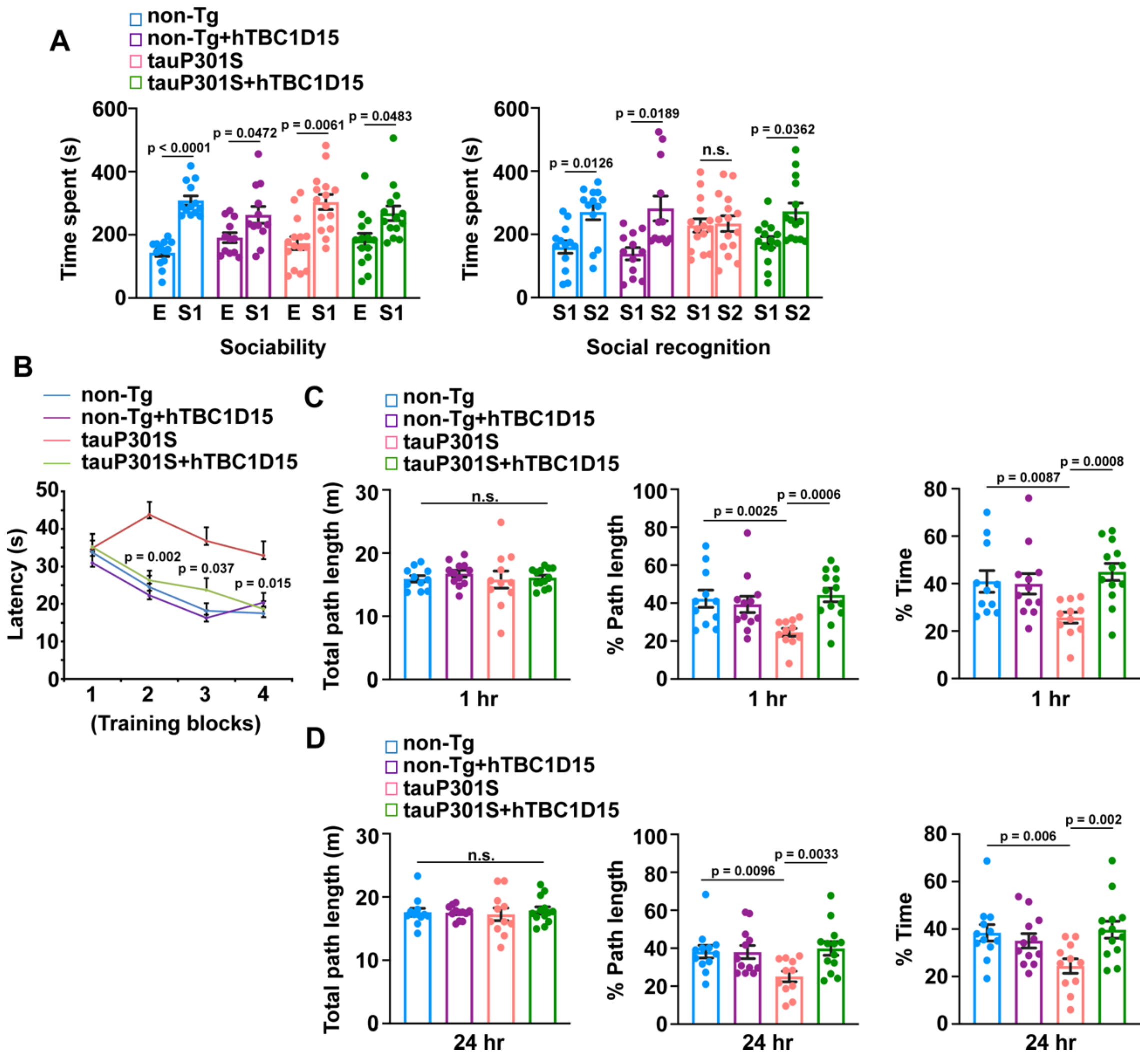


**Fig. S8.** TBC1D15 OE improves behavioral performance in tauopathy mice. (*A*) Sociability and social recognition memory assay performed in 6-month-old non-Tg and tauP301S Tg male mice injected with AAV-mCherry or AAV-mCherry-hTBC1D15 (*n* = 12-15 male mice per group). n.s.: non-significant. (*B-D*) Morris water maze test of 7-month-old tauP301S Tg male mice and non-Tg littermates with and without TBC1D15 OE (*n* = 11-13 male mice per group). Data were shown as the mean ± SEM and analyzed by two-way ANOVA, followed by two-sided paired Student's *t*-test (*A*) or one-way ANOVA, followed by Sidak's post-hoc test (*B-D*).

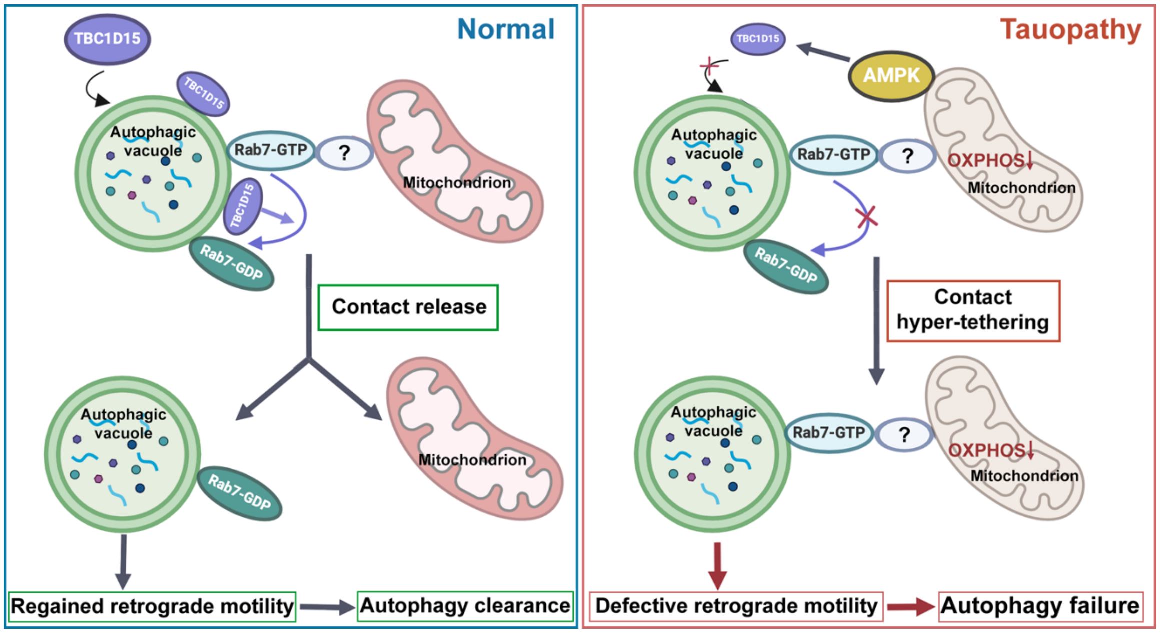


**Fig. S9.** A proposed model of a novel autophagosome/autophagic vacuole (AV)-mitochondria (Mito) contact dysregulation and the resulting autophagy defects in tauopathy neurons. Shown is a schematic illustration of a new class of inter-organelle interactions through AV-Mito membrane contacts in the axons of neurons. The contact dynamics of these two organelles is controlled by the contact release factor TBC1D15, a Rab7 GTPase-activating protein (GAP). In tauopathy neurons, mitochondrial bioenergetic dysfunction-induced hyperactivity of AMP-activated protein kinase (AMPK) accelerates TBC1D15 turnover, leading to TBC1D15 deficiency and excessive AV-Mito contact tethering. As a result, AV-Mito contact release defects curb AV retrograde transport and disrupt autophagic cargo clearance, thereby contributing to autophagy failure and pathological tau buildup in tauopathy.

### Tables

**Table S1.** Human brain specimens used for Western blot and qRT-PCR measurements

| Case type | Age/sex | Postmortem interval | Braak stage of AD/FTD brains |
|---|---|---|---|
| Control #1 | 55/M | 17.8 hr | 0 |
| Control #2 | 52/M | 19.55 hr | 0 |
| Control #3 | 62/M | 17.07 hr | 0 |
| Control #4 | 53/F | 15.98 hr | 0 |
| FTD #1 | 74/F | 19.08 hr | Braak VI |
| FTD #2 | 66/M | 7.25 hr | Braak VI |
| FTD #3 | 64/F | 15.83 hr | Braak I |
| FTD #4 | 65/M | 18.97 hr | Braak V |
| FTD #5 | 71/F | 16.2 hr | Braak III |
| FTD #6 | 76/F | 6.38 hr | Braak III |
